# The genomic basis of evolutionary novelties in a leafhopper

**DOI:** 10.1101/2022.06.28.497946

**Authors:** Zheng Li, Yiyuan Li, Allen Z. Xue, Vy Dang, V. Renee Holmes, J. Spencer Johnston, Jeffrey E. Barrick, Nancy A. Moran

## Abstract

Evolutionary innovations generate phenotypic and species diversity. Elucidating the genomic processes underlying such innovations is central to understanding biodiversity. In this study, we addressed the genomic basis of evolutionary novelties in the Glassy-Winged Sharpshooter (*Homalodisca vitripennis*, GWSS), an agricultural pest. Prominent evolutionary innovations in leafhoppers include brochosomes, proteinaceous structures that are excreted and used to coat the body, and obligate symbiotic associations with two bacterial types that reside within cytoplasm of distinctive cell types. Using PacBio long-read sequencing and Dovetail Omni-C technology, we generated a chromosome-level genome assembly for the GWSS, then validated the assembly using flow cytometry and karyotyping. Additional transcriptomic and proteomic data were used to identify novel genes that underlie brochosome production. We found that brochosome-associated genes include novel gene families that have diversified through tandem duplications. We also identified the locations of genes involved in interactions with bacterial symbionts. Ancestors of the GWSS acquired bacterial genes through horizontal gene transfer (HGT), and these genes appear to contribute to symbiont support. Using a phylogenomics approach, we inferred HGT sources and timing. We found that some HGT events date to the common ancestor of the hemipteran suborder Auchenorrhyncha, representing some of the oldest known examples of HGT in animals. Overall, we show that evolutionary novelties in leafhoppers are generated by the combination of acquiring novel genes, produced both *de novo* and through tandem duplication, acquiring new symbiotic associations that enable use of novel diets and niches, and recruiting foreign genes to support symbionts and enhance herbivory.

## Introduction

Evolutionary innovations generate phenotypic novelties, help organisms expand into new ecological niches, and often result in lineage diversification (Hunter 1998; Wagner and Lynch 2010; Rabosky 2017). Classic examples of evolutionary novelties include the wings of insects (Nicholson et al. 2014) and the flowers of angiosperms (Endress 2011). Multiple hypotheses have been forwarded for how evolutionary novelties originate and evolve (Wagner 2011). At the gene level, two major molecular mechanisms have been proposed. The first is based on modifications of existing genes. Multiple studies have shown that changes in the regulation of gene expression or in existing coding gene sequences can generate phenotypic novelties (Carroll 2008; Blount et al. 2012). The second category involves the acquisition of novel genes. New genes can be acquired from foreign organisms through horizontal gene transfer (HGT, or lateral gene transfer) (Moran and Jarvik 2010; Acuña et al. 2012; Wybouw et al. 2016; Husnik and McCutcheon 2018; McKenna et al. 2019; Xia et al. 2021), *de novo* gene birth from non-coding sequences (Long et al. 2013; Jin et al. 2021), and from gene or genome duplication followed by the divergence of paralogous genes (Ohno 1970; Lynch and Force 2000; Birchler and Yang 2022). In eukaryotes, gene duplication is the most common mechanism for creating new genes. At the cellular and organismal level, novel phenotypes and functional capabilities can be generated through the acquisition of symbionts (Perreau and Moran 2022). In turn, novel symbiotic relationships can result in new types of cells and organs, resulting in increased phenotypic complexity over time (Moran 2007; Perreau and Moran 2022).

The origin of evolutionary novelties can be complex and may involve different mechanisms at multiple levels of biological organization. In this study, we use a leafhopper called the Glassy-Winged Sharpshooter (*Homalodisca vitripennis*, GWSS) (Fig. 1A) as a study system to understand the genomic basis of several distinct types of evolutionary novelties. Leafhoppers (Insecta: Hemiptera: Cicadellidae) constitute the second largest hemipteran family with more than 20,000 species (Wahlberg et al. 2006).

**Fig 1.**
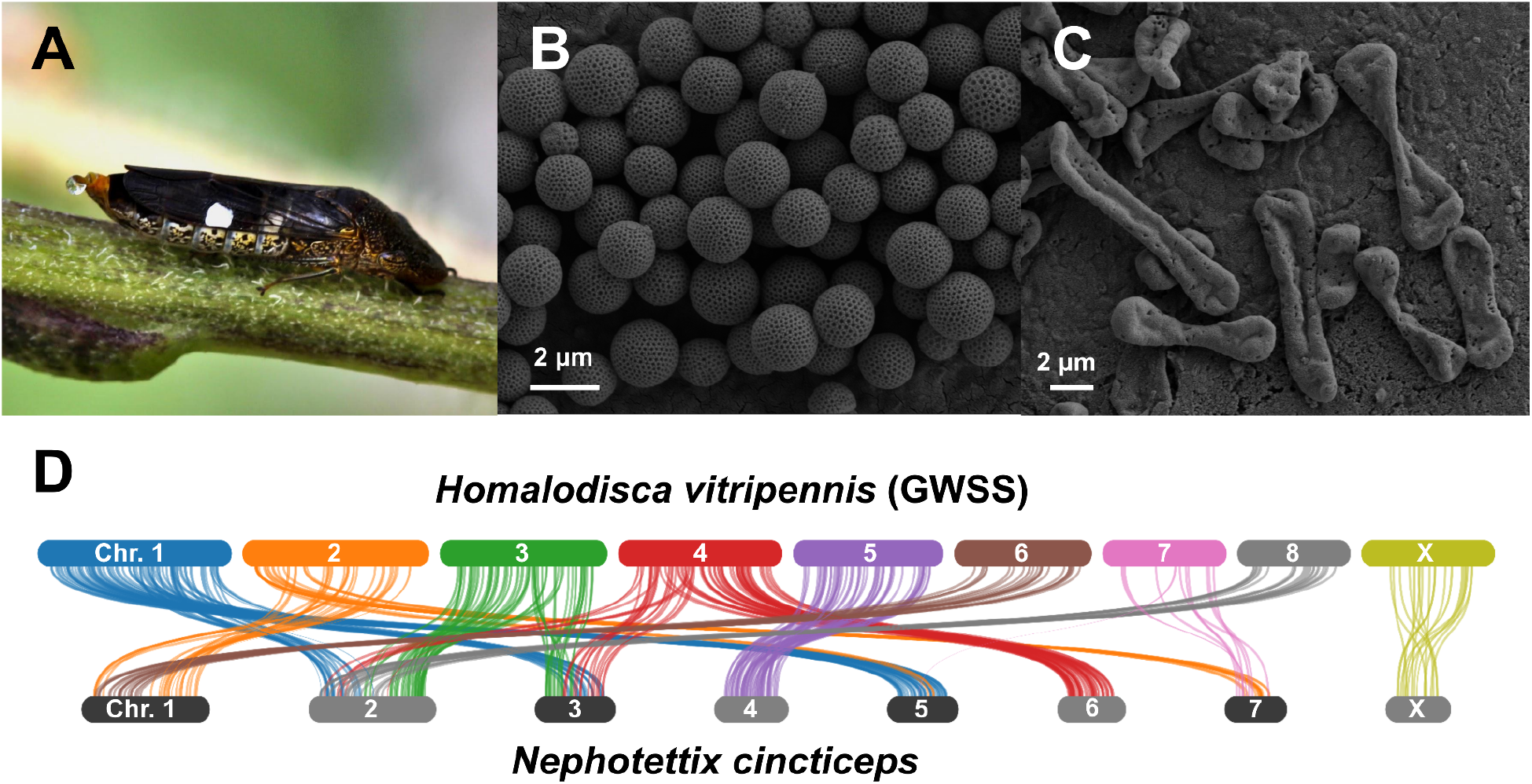
(**A**) *Homalodisca vitripennis* female (Glassy-Winged Sharpshooter, GWSS); (**B**) Scanning electron microscopy (SEM) of GWSS integumental brochosomes; (**C**) SEM of GWSS egg brochosomes; (**D**) Syntenies between two leafhoppers, GWSS vs. *Nephotettix cincticeps* (Rice Green Leafhopper). Bars represent chromosomes. The length of the bars is proportional to the length of the chromosome-level scaffolds in the assemblies.

One prominent evolutionary innovation in leafhoppers is the production of proteinaceous nanostructures called brochosomes (Fig. 1B, C). Brochosomes are secreted into the hindgut lumen by cells in a portion of the Malpighian tubules, then excreted. Leafhoppers then spread brochosomes using distinctive setae on their hind legs to coat their bodies and, in some species, egg masses (Tulloch and Shapiro 1954; Day and Briggs 1958; Rakitov 1999; Rakitov et al. 2018). Brochosomes are hydrophobic and prevent fouling of leafhopper surfaces by their sugary exudates, and it has also been hypothesized that they are involved in camouflage and preventing microbial infections (Rakitov 2004; Rakitov and Gorb 2013a; Rakitov and Gorb 2013b; Yang et al. 2017). Molecular phylogenies support a single origin of brochosomes in the ancestor of leafhoppers. Four novel gene families are involved in brochosome production (Rakitov et al. 2018). The largest of these families was reported to contain 28 paralogs within a single genome, suggesting gene duplication during the evolution of these novel leafhopper nanostructures.

Most leafhoppers feed on plant phloem or xylem sap, in which essential amino acids are scarce. These nutrients are supplied by two bacterial symbionts, *Sulcia* and *Nasuia*, which are among the oldest insect symbionts, acquired over 200 million years ago in a common ancestor of the hemipteran suborder Auchenorrhyncha (Moran et al. 2005; Bennett and Moran 2013; Cao and Dietrich 2022). These symbionts are hosted in specialized organs called bacteriomes. Interestingly, the sharpshooters (subfamily Cicadellinae) acquired a new bacterial symbiont, *Baumannia*, which replaced *Nasuia* (Moran et al. 2003; Takiya et al. 2006). This acquisition coincided with a shift from phloem-feeding to xylem-feeding and resulted in the generation of a novel cell type that hosts *Baumannia* (Moran 2007).

During the long-term evolution of intracellular bacterial symbionts, such as *Sulcia* and *Nasuia*, the symbiont genomes shrink over time due to relaxed selection and genetic drift causing the loss of non-essential genes (McCutcheon and Moran 2011; Bennett and Moran 2013; Bennett and Moran 2015). In several cases, sap-feeding insects have acquired foreign genes from other microbes through HGT; some of these acquired genes support interactions with symbionts (Mao et al. 2018; Van Leuven et al. 2019; Mao and Bennett 2020). Gene acquisitions by HGT can also facilitate other evolutionary novelties in these insects. For example, aphids acquired genes for carotenoid biosynthesis from fungi (Moran and Jarvik 2010; Nováková and Moran 2012). In whiteflies, genes transferred from plants enable the detoxification of plant compounds (Xia et al. 2021). In leafhoppers and cicadas, multiple HGT events have been documented for genes that appear to support interactions with symbionts or to enhance herbivory (Husnik and McCutcheon 2018; Mao et al. 2018; Van Leuven et al. 2019; Mao and Bennett 2020). Thus, HGT appears to be a central mechanism by which sap-feeding insects have colonized new ecological niches and diversified.

In this study, we applied PacBio long-read sequencing and the Dovetail Omni-C technique to generate a chromosome-level genome assembly for the GWSS, a sap-feeding insect that is one of the most-studied leafhopper species because it is an agricultural pest. We evaluated the quality and completeness of the assembly using karyotyping and flow cytometry. By combining this assembly with transcriptomic and proteomic data, we investigated the genomic basis of three evolutionary novelties: (1) novel proteins underlying the production of brochosomes; (2) interactions with a new symbiont, *Baumannia*; (3) foreign gene acquisitions by HGT. We also inferred the phylogenetic placement of these HGT events. Our study reveals that a combination of different genetic mechanisms contributed to the evolutionary novelties that fueled leafhopper diversification.

## Results

### Assembly and annotation of the GWSS genome

We assembled the GWSS genome from 237.9 Gb of Pacbio long reads and 53.8 Gb of Omni-C reads. After genome polishing, our assembly had an overall length of 2.3 Gb with a scaffold N50 of 168.8 Mb. Nine scaffolds were larger than 132.9 Mb, and suggested a haploid chromosome count of n=9. We found no published karyotype information for GWSS, so we performed karyotyping on males to estimate the chromosome number and to identify the X chromosome. We found a haploid chromosome number of n=9 in the GWSS, which is consistent with the nine large scaffolds from our genome assembly (Fig. S1, Table 1). The total length of the nine chromosome-level scaffolds is 1.7 Gb, which is 74.1% of the total genome assembly (Table 1, Table S1).

**Table 1.**
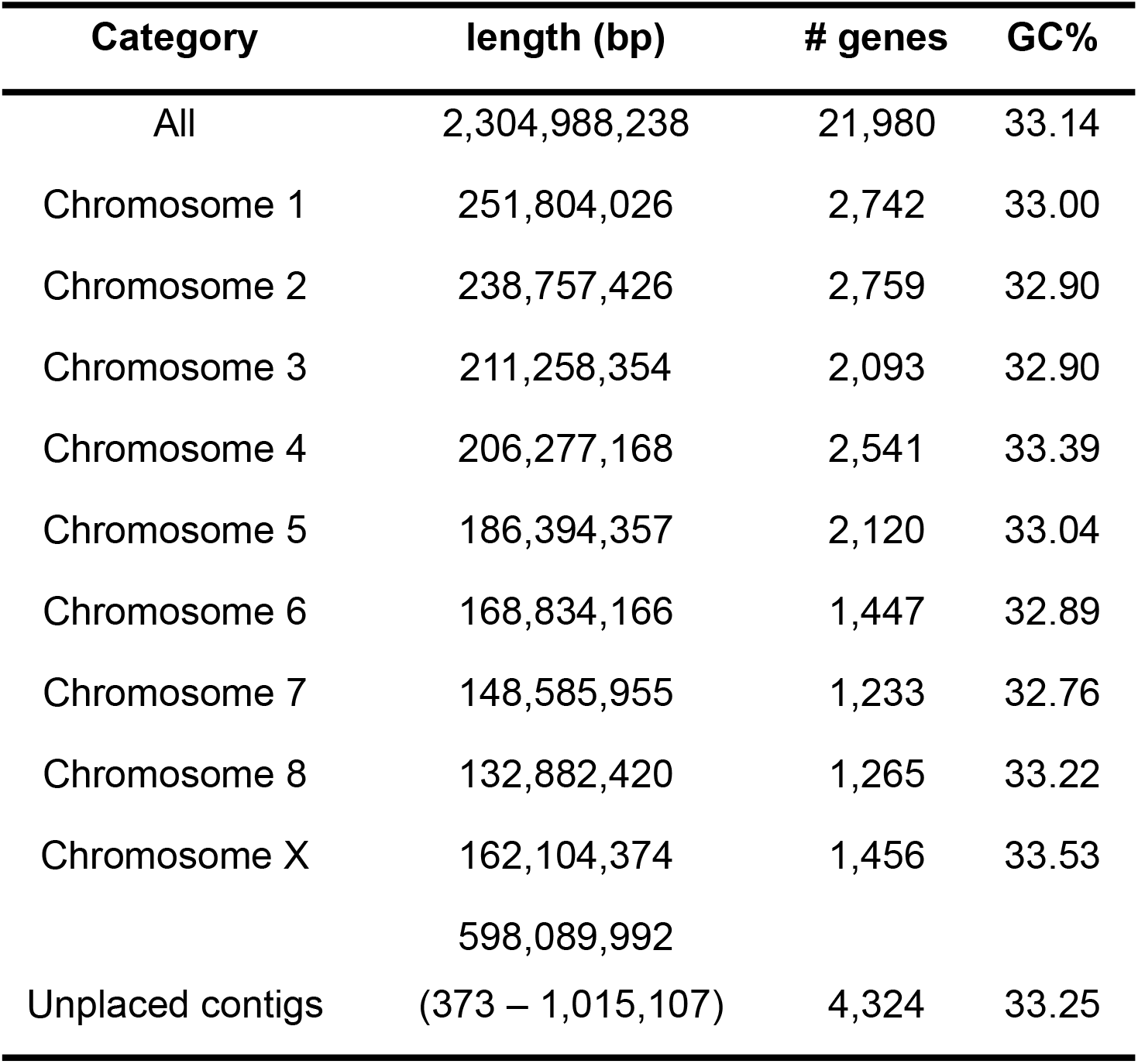
Summary of the chromosome-level genome assembly of the Glassy-Winged Sharpshooter.

Most leafhoppers have an XO sex determination system (Tree of Sex Consortium 2014). To identify the X chromosome in the GWSS genome, we mapped male and female Illumina reads to our nine chromosome-level scaffolds. Given the XO sex determination system, we expect the male vs. female sequencing depth of the X chromosome to be approximately half what it is for the autosomes. In our genome assembly, eight scaffolds had similar normalized read depths between sexes. By contrast, the seventh largest scaffold (162.1 Mb) had about half of the normalized sequencing read depth ratio between sexes when compared to other chromosomes (Fig. S2). Therefore, we inferred that the seventh-largest scaffold to be the X chromosome. We subsequently named the other eight autosomes as chromosomes 1-8 based on their sizes, in order from longest to shortest (Table 1).

To further evaluate our assembly, we estimated genome size using flow cytometry. This provided an estimate of 1.89 Gb and 1.99 Gb for male and female genome sizes, respectively (Table S2). The X chromosome size estimated by flow cytometry (199 ± 40 Mb) is consistent with the inferred X chromosome size (162.1 Mb), further supporting its assignment as the sex chromosome. The DNA content estimated by flow cytometry is also consistent with the total length of the nine chromosome-level scaffolds, supporting the overall accuracy of our GWSS genome assembly.

We used Benchmarking Universal Single-Copy Orthologs (BUSCO) to evaluate the completeness of the genome assembly (Simão et al. 2015). Querying the single copy orthologs of Hemiptera resulted in a BUSCO score for the GWSS assembly of 93.7% (92.7% single and duplicated, 1.0% fragmented, 6.2% missing). This score is similar to that of the other recent GWSS genome assembly (Ettinger et al. 2021). The BUSCO score for the nine chromosome-level scaffolds alone was 90.8% (86.6% single and duplicated, 4.2% fragmented, 9.2% missing)(Table S3).

The NCBI RefSeq annotation pipeline was used to annotate the genome (O’Leary et al. 2016). We used WindowMasker (Morgulis et al. 2006) to mask repetitive elements, which constituted 39.4% of the genome. We then aligned 42 GWSS transcriptomes containing 3,410,135,668 reads onto the repeat-masked genome. We predicted a total of 21,980 annotated genes with 19,904 protein-coding genes and 611 pseudogenes. 17,656 annotated genes were found on the nine chromosome-level scaffolds, with 1,456 on the X chromosome and 16,200 on the autosomes (Table 1, Table S1).

### Comparative genomics of leafhoppers and planthoppers

To compare the genome content and organization of leafhoppers (Hemiptera: Auchenorrhyncha: Cicadellidae) and planthoppers (Hemiptera: Auchenorrhyncha: Fulgoroidea), we used the three available chromosome-level genome assemblies, including the GWSS, *Nephotettix cincticeps* (rice green leafhopper), and *Nilaparvata lugens* (brown planthopper). Based on gene cluster assessment, 6,989 gene families are shared by these three species. The two leafhoppers share 2,337 gene clusters that are not found in *Ni. lugens* (Fig. S3). The GWSS and *Ne. cincticeps* shared 519 gene families that are absent in *Ni. lugens*. We also compared synteny between these three species. We observed little synteny between GWSS or *Ne. cincticeps* when aligned to *Ni. lugens* (Fig. S4). We found 113 syntenic blocks between the two leafhoppers (Fig. 1D). The X chromosome exhibited similar conservation of gene content and arrangement between species to what was found for the autosomes.

### Genomic distribution of brochosome-related genes

Brochosomes are produced by specialized glandular segments of the Malpighian tubules in leafhoppers (Rakitov et al. 2018). A previous study using transcriptomics and proteomics identified four major gene families related to brochosome production; these families are referred to as brochosomins, glycine-rich, poly-proline, and cyclase-like proteins (Rakitov et al. 2018). However, the molecular and genetic basis of brochosome production is still not fully understood. We used this list of brochosome-related genes to locate homologous sequences in the GWSS genome. Overall, we identified 68 brochosome-related genes from these four major gene families.

We next analyzed the locations of each brochosome-related gene in the GWSS genome (Fig. 2, Table S4). We found 28 genes in the brochosomin gene family. Of these, 21 are located in two major clusters on the X chromosome (Fig. 2, Table S4). We also observed 33 genes in the glycine-rich family, distributed across multiple chromosomes, with a cluster of 21 genes on chromosome 7. We found one poly-proline gene and five cyclase-like genes on autosomes.

**Fig 2.**
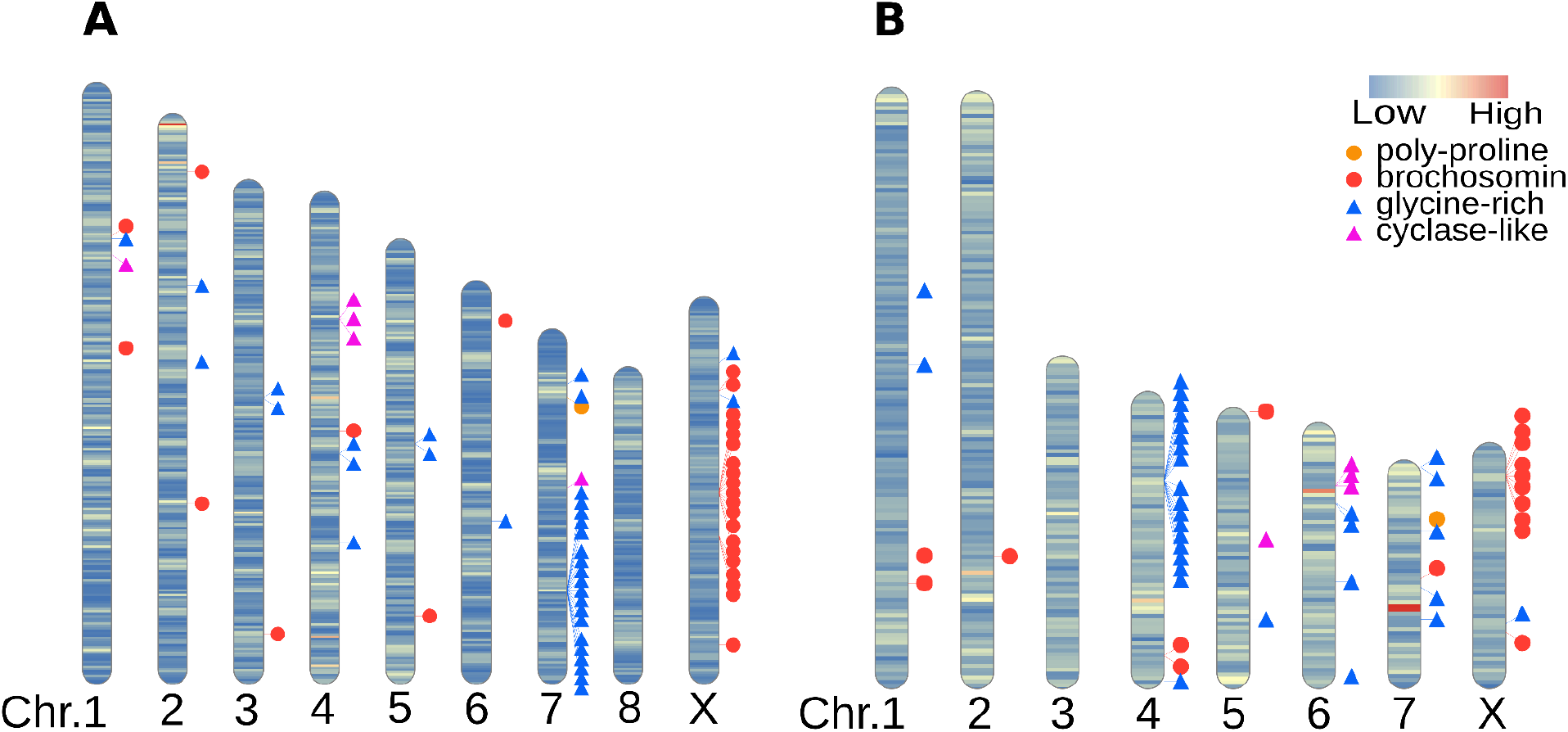
Ideogram with chromosomal locations of brochosome related genes in two leafhopper genomes. The intensity of the color on each chromosome represents gene density. (**A**) GWSS genome. (**B**) *Nephotettix cincticeps* (rice green leafhopper) genome.

To compare repertoires of brochosome-related genes between leafhopper species, we used the reciprocal best blast hits of the GWSS brochosome-related genes to locate homologous sequences in the *Ne. cincticeps* genome. We found 53 brochosome-related genes from the four gene families in the rice green leafhopper genome (Fig. 2, Table S4). We found only 17 genes in the brochosomin gene family for *Ne. cincticeps*, suggesting that the GWSS has undergone more duplications of these loci. Nine brochosomin genes form a cluster on the likely X chromosome based on its homology to the X chromosome of the GWSS. We also found a gene cluster of 17 genes in the glycine-rich family on chromosome four. As for the GWSS, we observe one poly-proline gene and five cyclase-like genes on autosomes of *Nephotettix*.

### Expression levels of brochosome-related genes

To examine the expression of brochosome-related genes, we sequenced transcriptomes from the GWSS Malpighian tubules and control samples that consisted of pooled tissues from the gut and other internal organs from the same individuals, with three biological replicates of each type of sample. We mapped transcriptome reads to the genome and performed differential expression analyses. We found that 1,190 genes are significantly upregulated in the Malpighian tubules (Table S5). Given that Malpighian tubules have other biological functions, unrelated to brochosome production, we focused on the expression of known brochosome-related genes. We found that 37 of the 68 identified brochosome-related genes are upregulated in the Malpighian tubules (Fig. 3). The other 31 that do not show this pattern might be expressed at different times or have functions unrelated to brochosome production.

**Fig 3.**
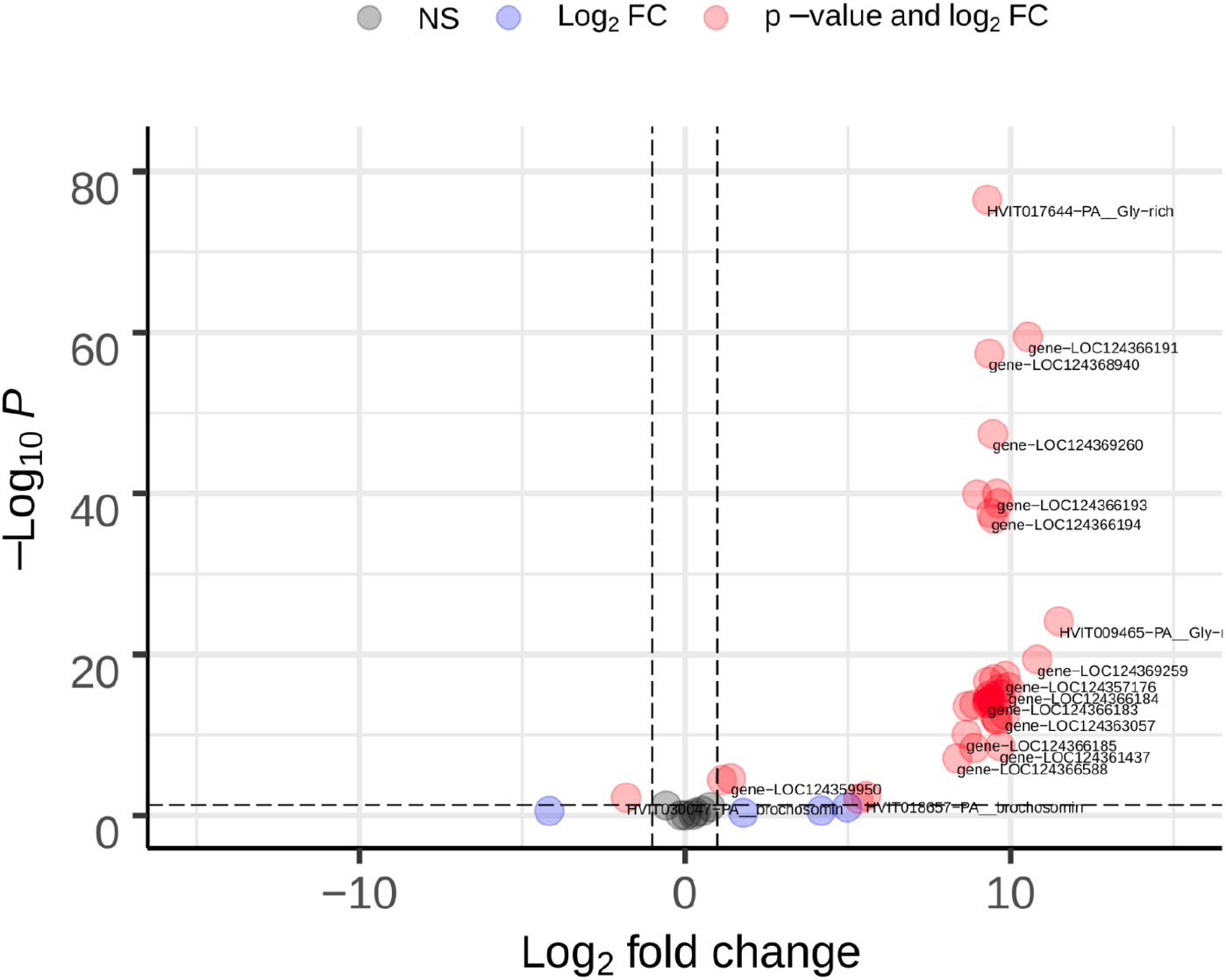
Differential expression of 68 brochosome-related genes in Malpighian tubules vs. other internal organs (n Malpighian tubules=3, other internal organs=3). Horizontal dashed line: cut-off for the adjusted p-value < 0.05. Vertical dashed lines: Log_2_ fold change > |1|. Differentially expressed brochosome-related genes exceeding both cut-offs are shown in red. Genes exceeding only the Log_2_ fold change cut-off are shown in blue. Genes that are not significantly differentially expressed are shown in black.

To examine the expression of brochosome-related genes at the protein level, we used LC-MS/MS to compare with the proteins found in the Malpighian tubules and the rest of the gut from the GWSS abdomen. We detected 3,537 proteins across all proteomics samples. We found that 13 brochosome-related proteins are significantly upregulated in Malpighian tubules vs. the controls (Table S6). Seven of these were also upregulated in the transcriptome analyses. We also found a cyclase-like protein that is detected only in the pooled tissues that include the gut. In agreement with this observation, the gene was not significantly upregulated in the Malpighian tubule transcriptome. To determine which of these proteins are also found in brochosomes, we performed LC-MS/MS proteomics on three samples containing brochosomes and other biomolecules isolated from the insect integument. We identified eleven brochosome-related proteins and five of these are among the top 150 most abundant proteins in terms of the peptide-spectrum matches (PSMs) in these samples. Seven of these eleven proteins found in brochosomes were also found in the Malpighian tubule proteomics samples and were significantly upregulated in the Malpighian tubule transcriptomes (Fig. 3, Table S7).

### Symbiosis-related genes in the GWSS

The GWSS has two types of bacteriomes for hosting endosymbionts. The red bacteriome contains a relatively recently (∼ 30 Mya) acquired symbiotic bacterium, *Baumannia*. The yellow bacteriome hosts *Baumannia* and *Sulcia*, an older (∼250 Mya) symbiont that is found in most leafhoppers as well as most other Auchenorrhyncha. A previous study identified host genes that are upregulated in these two bacteriomes using transcriptome-based assemblies as the reference (Mao and Bennett 2020). To re-evaluate their differential expression analyses based on the new reference genome, we mapped this RNAseq data from this study to the GWSS genome. In the red bacteriome, 1,504 host genes are upregulated, and, in the yellow bacteriome, 2,063 genes are upregulated. Overall, 521 host genes are upregulated in both bacteriomes (Table S8), which is significantly more overlap between the two sets of genes than expected by chance (hypergeometric test, p < 10^−273^). We also mapped the genes upregulated in each type of bacteriome to our assembled chromosomes. The overlapping high gene density regions show that some chromosomal locations of upregulated genes are shared between the two bacteriomes (Fig. S5).

### Evolutionary history of HGT genes

Previous studies have revealed genes arising from horizontal gene transfer (HGT) in leafhoppers, with at least fourteen such genes in the GWSS (Mao et al. 2018; Mao and Bennett 2020). To study the evolutionary history of these HGT genes, we first used the GWSS genome to identify their chromosomal locations. We observed that the majority of HGT genes are located on autosomes, with six on chromosome 1 and two on chromosome 2. One HGT gene, the *ATPase* gene, is found on the X chromosome (Fig. S6). We also confirmed that there are multiple paralogs of the horizontally acquired *cel, gh25*, and *plc* genes. The paralogs of *cel* and *gh25* were found on the same chromosomes (Fig. S6, Table S9).

To estimate the phylogenetic placement of the HGT sources, we assembled a dataset with 102 transcriptomes and 25 genomes across the Auchenorrhyncha (Table S10). We identified sequences that were homologous to the fourteen HGT genes in this dataset and constructed gene family phylogenies for these genes (Fig. 4, Fig. S7). Based on gene presence and absence in the context of the gene tree topologies, our results support multiple independent HGT events from different bacterial sources to leafhoppers. Strikingly, the *pel* genes were present in Cercopoidea, Cicadoidea, and Membracoidea, and the gene tree topology supports a single origin through HGT in the common ancestor of these three superfamilies, which span the Auchenorrhyncha suborder. We also found that the *alv* gene was likely horizontally transferred to the common ancestor of Auchenorrhyncha (Fig. 4, Fig. S7, Table S9). Similarly, *cel, def, gh25*, and the other four genes were possibly transferred to the common ancestor of Membracoidea. In contrast to these HGT genes with deep origins, the *per* and *rluA* genes were only found in one or two species of leafhoppers, suggesting relatively recent HGT events (Fig. 4, Fig. S7, Table S9). We found orthologs of the *plc* gene in the Coleorrhyncha and Auchenorrhyncha, but the phylogeny of the gene tree suggests two independent transfers of *plc* in these two suborders (Fig. 4, Fig. S7, Table S9).

**Fig 4.**
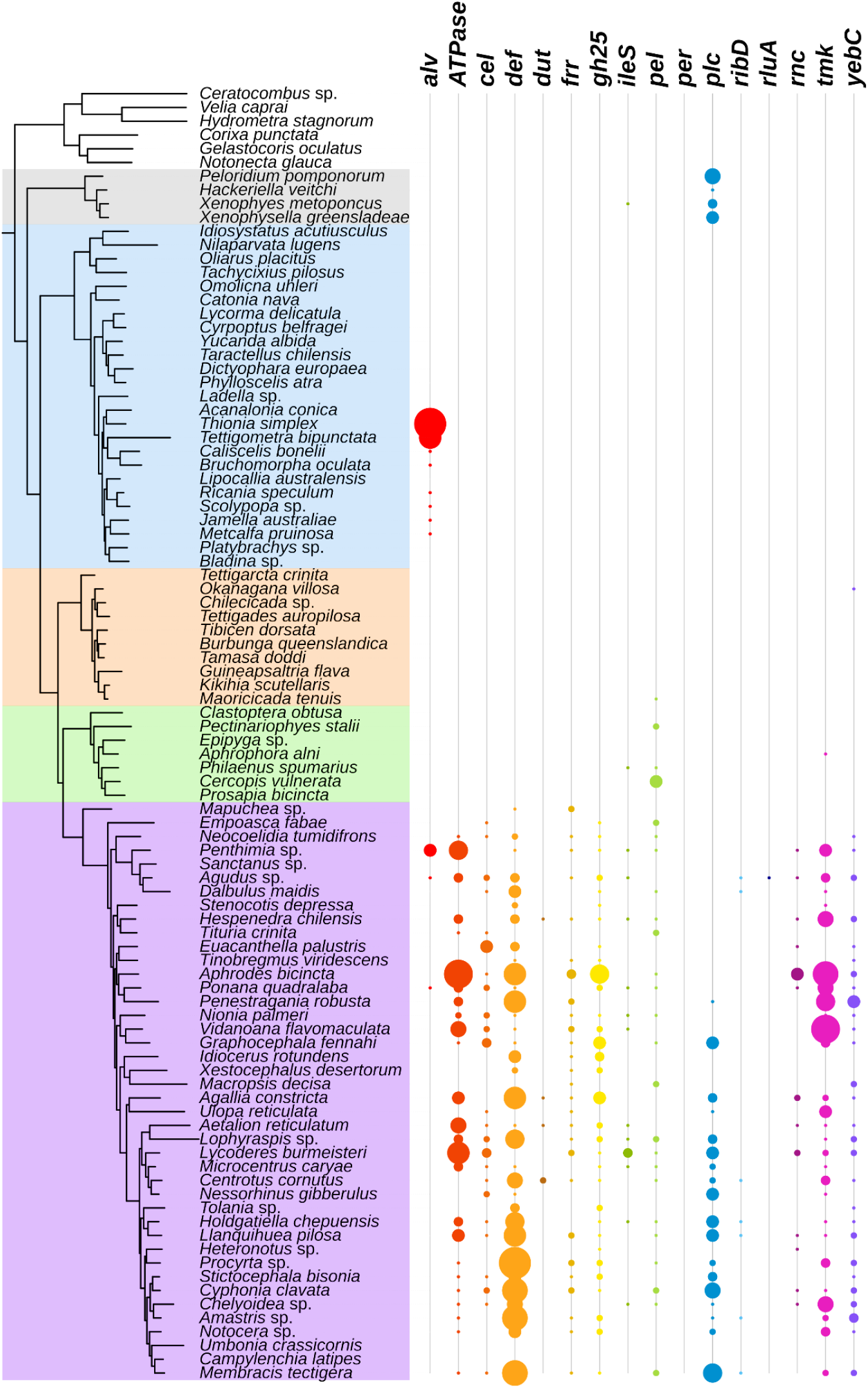
Phylogeny of Auchenorrhyncha and HGT gene copy numbers in transcriptomes. The phylogeny is adopted from Skinner *et al*. 2020, the colors correspond to different major lineages (White: Outgroups; Gray: Coleorrhyncha; Blue: Fulgoridae; Orange: Cicadoidea; Green: Cercopoidea; Purple: Membracoidea). The size of each circle represents the gene copy number found in a given transcriptome.

## Discussion

### Genome assembly of the GWSS

Leafhoppers are a highly diverse insect lineage and include some major agricultural pest species. The GWSS itself is a significant invasive pest species, primarily as a vector of bacterial pathogens, including *Xylella fastidiosa* in grapes and other woody hosts. To date, the only other chromosome-level genome assembly for a leafhopper is that for *Neophotettix cincticeps* in the subfamily Deltocephalinae (Yan et al. 2021). Our GWSS genome represents the subfamily Cicadellinae, which diverged from Deltocephalinae over 125 million years ago (Cao et al. 2022). Our genome N50 is higher than a recent assembly for GWSS that was based on PacBio and Nanopore reads (Ettinger et al. 2021). Our total assembly length is larger but consistent with the other assembly. Karyotyping confirmed that the chromosome number corresponds to the number of large scaffolds in our genome assembly. Furthermore, the combined length of the chromosome-level scaffolds is close to the genome size estimated from flow cytometry. Thus, the many small scaffolds likely represent a combination of contaminants plus sequences on homologous chromosomes that did not assemble due to heterozygosity in our sample. Overall, the agreement between the assembly and empirical approaches suggests we have a high-quality and accurate genome.

### Comparative genomics of Auchenorrhyncha

We identified and compared syntenic regions between the two leafhopper genomes. The patterns suggest chromosome fission/fusion occurred during the evolution of these species. Little synteny was found between leafhoppers and planthoppers. This result is not surprising given the deep divergence between these two lineages. Recent genomic studies of aphids have found that gene content and arrangements are highly conserved on the X chromosome as compared to autosomes (Li et al. 2020; Mathers et al. 2021), but leafhoppers did not show higher conservation of the X chromosomes. In this regard, leafhoppers resemble other hemipteran insects such as kissing bugs and planthoppers, in which the X is also not more conserved than autosomes (Mathers et al. 2021). This suggests that the pattern is unique to aphids and reflects their unusual reproductive biology (Li et al. 2020; Mathers et al. 2021).

### Brochosome and Brochosome-related genes

Brochosomes are novel proteinaceous structures found only in leafhoppers (Tulloch and Shapiro 1954; Day and Briggs 1958; Rakitov 1999; Rakitov et al. 2018). Four major gene families have been hypothesized to underlie brochosome production (Rakitov et al. 2018). A previous study based on transcriptomes revealed multiple copies of the brochosomin, glycine-rich, and cyclase-like gene families per genome (Rakitov et al. 2018). We found that many of these genes were organized into gene clusters. In GWSS and *Nephotettix*, there are two and one major gene clusters of brochosomin genes, respectively. In GWSS, the glycine-rich genes form a cluster on chromosome 7. Similarly, glycine-rich genes also form a cluster on chromosome 4 of *Nephotettix*. No synteny is found between chromosome 7 of the GWSS and chromosome 4 of *Nephotettix*. This observation suggests that glycine-rich genes expanded independently by tandem duplication in these two leafhopper lineages. In both genomes, we observed that cyclase-like genes have tandem duplicates on an autosome. Overall, we found that tandem duplication plays an important role in the expansion and diversification of these novel gene families. Unlike these cases, we confirmed that the remaining brochosome-associated gene family, encoding the poly-proline protein, exists as a single copy in both the GWSS and *Nephotettix* genomes.

The two brochosomin gene clusters of the GWSS are situated on the X chromosome, and the single brochosomin gene cluster of *Nephotettix* is on what is probably the X chromosome, based on its homology to the GWSS X. Interestingly, it has been shown that some sharpshooters, including GWSS, produce two kinds of brochosomes (Rakitov 2004). These include the spherical integumental brochosomes that most leafhoppers excrete and anoint onto their integument, and the so-called egg brochosomes, which are larger and more elongated (Rakitov 2000). Egg brochosomes, produced only by females, are collected as a deposit on the wings and then used to cover the egg masses following oviposition into leaf tissue (Rakitov 2004). *Nephotettix* belongs to the Deltocephalinae, which only produces integumental brochosomes (Rakitov 2004). Potentially, the differences in brochosomin gene numbers and brochosomin gene clusters on the X chromosome are related to the female-specific production of egg brochosomes in GWSS.

In addition to genomic analyses, we used transcriptome analyses and shotgun proteomics to identify candidate genes that are important for brochosome production. Transcriptomics and shotgun proteomics of the Malpighian tubules identified 37 and 13 genes from brochosome-related gene families that are highly upregulated, respectively. We detected eleven of these proteins in proteomic analyses of partially purified brochosomes. This evidence from both Malpighian tubules and brochosomes themselves supports the importance of these genes for brochosome production. By cross-referencing genomes, transcriptomes, and proteomics, our study provides a comprehensive candidate list of brochosome-related genes of the GWSS.

Previous studies have found tremendous morphological diversity in brochosomes (Rakitov 2004; Rakitov et al. 2018). For example, the diameters of the regular spherical brochosomes produced by most species vary from 200 to 700 nm but may be as large as 5 μm (Rakitov 2004). It remains to be understood what genes and proteins are responsible for this variation and whether it has any significance in terms of insect fitness. Potentially, gene duplication has contributed to this morphological diversity. Future experiments could address the biological consequences of silencing brochosome-related genes. CRISPR/Cas9 genome editing was recently demonstrated in GWSS (de Souza Pacheco et al. 2022), and this approach might be applied to test how disrupting genes we identified impacts brochosome biogenesis and morphology. Our high quality GWSS genome and the comprehensive list of brochosome-related genes will serve as a foundation for understanding the biology of brochosomes.

### Symbiosis-related genes in GWSS

Sharpshooters, including the GWSS, acquired a novel symbiont, *Baumannia* (Moran et al. 2003; Takiya et al. 2006). The red bacteriome of the GWSS represents a novel organ that supports this new symbiosis, and it has a distinctive pattern of gene expression (Mao and Bennett 2020). By mapping transcriptome reads of both bacteriome types to our genome assembly, we evaluated symbiosis-related genes potentially used to support each endosymbiont. Consistent with the previous study, we found more upregulated genes in the more ancient yellow bacteriome, which hosts both *Baumannia* and *Sulcia*, potentially linked to the fact that the *Sulcia* genome is more highly reduced. The new chromosome-level genome assembly allowed us to identify the locations of symbiosis-related genes, as was done in previous analyses for aphids and a psyllid, in suborder Sternorrhyncha (Li et al. 2020). We found that symbiosis-related genes occur on all GWSS chromosomes. We did not observe an enrichment of these genes on the autosomes relative to the X chromosome, as is observed in aphids (Li et al. 2020). In the GWSS genome, a similar overall profile of chromosomal regions supports each bacteriome. In part, this reflects the significant overlap in the sets of GWSS genes that are upregulated in each type of bacteriome. This pattern is consistent with previous observations (Mao and Bennett 2020).

### Horizontal gene transfers of Auchenorrhyncha

During their evolution, insects have occasionally acquired genes from bacteria, fungi, and plants (Moran and Jarvik 2010; Acuña et al. 2012; Wybouw et al. 2016; Husnik and McCutcheon 2018; McKenna et al. 2019; Xia et al. 2021). Previous studies found multiple HGT genes in draft genomes of the GWSS and other leafhoppers (Mao et al.2018; Mao and Bennett 2020). We confirmed the presence of these genes in our genome assembly for GWSS and identified their chromosomal locations. We found that the majority of the HGT genes are on the autosomes of the GWSS; only the ATPase gene is located on the X chromosome. Paralogs of *cel* and *gh25* are found as tight clusters on the same chromosomes. These observations suggest that these two HGT genes have been duplicated by tandem duplications.

Previous studies have shown that some HGT genes are found in lineages of leafhoppers and cicadas, suggesting a common ancestry of these genes in these hemipteran insects (Mao et al. 2018; Van Leuven et al. 2019; Mao and Bennett 2020). However, the phylogenetic placement of the HGT events remained unresolved. To better place these HGT sources, we used a phylogenomic approach to study HGT gene presence and absence, and to infer HGT gene tree topologies across Auchenorrhyncha. We found that some HGT genes likely have deep origins. For example, the *alv* gene was likely transferred to the common ancestor of the Auchenorrhyncha, which places the time of this event at around 250-300 MYA (Johnson et al. 2018; Cao and Dietrich 2022), and possibly lost in the Cicadoidea (cicadas), Cercopoidea (spittlebugs), and Membracidae (treehoppers). Similarly, the *pel* genes are shared by the Cicadoidea, Cercopoidea, and Membracoidea. Based on molecular dating of the species phylogeny (Johnson et al. 2018), *pel* was acquired by the common ancestor of these major lineages around 250 MYA. These HGT genes are some of the oldest known examples of HGT in animals (Husnik and McCutcheon 2018). In contrast, *per* and *rluA* are only found within the Deltocephalinae, suggesting relatively recent HGT events.

Interestingly, we found the *plc* gene in moss bugs (Coleorrhyncha) and leafhoppers. The gene tree topology supports independent acquisitions of *plc* in these two lineages. Our gene tree supports an origin from *Providencia* in the Membracoidea (Mao et al. 2018; Mao and Bennett 2020), whereas moss bugs possibly acquired *plc* from a different bacterium. The *plc* genes have functions related to lipid metabolism, and they are not highly expressed in the bacteriomes of GWSS (Mao et al. 2018; Mao and Bennett 2020). The function and expression pattern of the *plc* genes in Coleorrhyncha is unknown. We identified different numbers of paralogs of HGT genes among closely related species. Although some of the variation may be due to the incomplete nature of transcriptomes, some is likely due to different levels of gene duplication following the HGT event. Overall, our study shows repeated HGT-based acquisitions of foreign genes during the evolution of Auchenorrhyncha.

## Conclusion

The agreement between the sequence assembly and empirical approaches supports the high-quality and accuracy of the GWSS genome assembly reported here. This genome and associated analyses illuminate the genetic underpinnings of different evolutionary novelties in the GWSS. In particular, our chromosome-level genome assembly will serve as a foundation for understanding the biology of leafhoppers and future studies of brochosomes.

## Materials and Methods

### Sample preparation for genome sequencing

Multiple *H. vitripennis* individuals were collected from crape myrtles (*Lagerstroemia indica*) near the campus of the University of Texas at Austin. For a high-quality genome assembly, a total of 0.6 g of male and female adult individuals were frozen and shipped to Dovetail Genomics (Santa Cruz, CA, USA). Three male individuals were used for DNA extraction and Pacbio long-read sequencing. Other male and female individuals were pooled for Dovetail Omni-C library preparation. The library was sequenced on an Illumina HiSeqX platform to produce approximately 30x sequence coverage.

### Assembly of the *H. vitripennis* genome

To assemble the *H. vitripennis* genome, 237.9 Gb of PacBio CLR reads were used as an input to WTDBG2 v2.5 (Ruan and Li 2020), minimum read length 20000, and minimum alignment length 8192. Additionally, realignment was enabled with the ‘-R’ option, and read type was set with the option ‘-x sq’. Blast results of the WTDBG2 output assembly against the NCBI nt database were used as input for blobtools v1.1.1 (Laetsch and Blaxter 2017), and scaffolds identified as possible contamination were removed from the assembly. Finally, purge_dups v1.2.3 (Guan et al. 2020) was used to remove haplotypic duplications.

The *de novo* genome assembly and Dovetail OmniC library reads were used as input data for genome scaffolding using the HiRise assembler version v2.1.6-072ca03871cc (Putnam et al. 2016). Dovetail OmniC library sequences were aligned to the draft input assembly using bwa (Li and Durbin 2009). The separations of Dovetail OmniC read pairs mapped within draft scaffolds were analyzed by HiRise to produce a likelihood model for the genomic distance between read pairs, and the model was used to identify and break putative misjoins, score prospective joins, and make joins above a threshold. The HiRise scaffolds were then polished by Nextpolish (Hu et al. 2020) using the PacBio long reads and OmniC library short reads used in the genome assembly and scaffolding.

To evaluate the completeness of our genome assembly, BUSCO version 3.0.2 (Simão et al. 2015) was used on the chromosome-level assembly using the single-copy orthologous gene set for Hemiptera from OrthoDB version 9 (Zdobnov et al. 2017).

### Genome annotation

The NCBI Eukaryotic Genome Annotation Pipeline was used for genome annotation (O’Leary et al. 2016). Repeat families found in the genome assemblies of *Homalodisca vitripennis* were identified and masked using WindowMasker (Morgulis et al. 2006). Over 20,000 transcripts of GWSS and high-quality proteins of GWSS and other closely related insects were retrieved from Entrez, aligned to the genome by Splign (Kapustin et al. 2008), minimap2 (Li 2018), or ProSplign (https://www.ncbi.nlm.nih.gov/sutils/static/prosplign/prosplign.html). Additionally, 3,410,135,668 reads from 42 GWSS RNA-Seq datasets were also aligned to the repeat-masked genome. Protein, transcript, and RNA-Seq read alignments were passed to Gnomon for gene prediction. Gnomon predictions selected for the final annotation set were assigned to models based on known and curated RefSeq and models based on Gnomon predictions. The overall quality of the annotations was assessed using BUSCO v4 (Seppey et al. 2019). The detailed annotation pipeline can be found at https://www.ncbi.nlm.nih.gov/genome/annotation_euk/process/.

### Chromosome number confirmation by karyotyping

Three male adult individuals of *H. vitripennis* were collected on the campus of the University of Texas at Austin. For karyotyping, the insects were injected with 50 μL of a 2% colchicine solution in the abdomen and left at room temperature overnight. They were then dissected in 1X PBS solution to separate and remove both testicular follicles from the upper abdomen. Testicular follicles were each transferred to 100 μL of 0.075 M sodium citrate solution for 10 min and subsequently fixed in modified Carnoy’s solution (3:1 absolute ethanol:glacial acetic acid) for 1 hour. Finally, they were each added to 100 μL of 50% acetic acid and the tissue was homogenized by blowing air into the solution with a micropipette. We then spotted 10 μL of the acetic acid solution containing testicular follicle cells onto slides and allowed it to air-dry at room temperature. All samples were stained with 15 μL Giemsa stain (5%) for 30 min, then rinsed completely and mounted in deionized water. The slides were viewed under a Nikon Eclipse te2000-u inverted fluorescence microscope, and cells with clear chromosome segregation were recorded and photographed with a Nikon DS-Ri2 Microscope Camera.

### Genome size estimation by flow cytometry

Genome size was estimated as described in (Johnston et al. 2019). In brief, a *H. vitripennis* head with unknown genome size and a *Drosophila virilis* female head standard (1C = 328 Mbp) were placed together into 1 mL of ice-cold Galbraith buffer in a 2 mL Dounce tissue grinder. Nuclei were released from both tissues by grounding with 15 gentle strokes using an A (loose) pestle, then filtered through nylon mesh into a 1.5 mL microfuge tube, and stained with 25 μl of propidium iodide (1 mg/ml) for 2 hours in the dark at 4 °C. The mean red PI fluorescence from 2C sample and standard nuclei was quantified using a CytoFlex flow cytometer (Beckman Coulter). Haploid (1C) DNA quantity was calculated as (2C sample mean fluorescence/2C standard mean fluorescence) × 328 Mbp. The X chromosome genome size was estimated as the genome size difference between XX females and XO males.

### Assignment of the X chromosome and autosomes

The X chromosome was assigned following the method previously used in the pea aphid and psyllid genomes (Li et al. 2019; Li et al. 2020). We mapped whole genome sequencing reads from male and female individuals back to our chromosome-level genome assembly. The male and female sequencing reads were obtained from the i5K insects genome project (Thomas et al. 2020) through GenBank (BioProject: PRJNA168119, Accession: SRX326930, SRX326929, SRX326928, SRX326927) and were cleaned with Trimmomatic version 0.38 (Bolger et al. 2014). The clean reads were mapped to the chromosome-level assembly using Bowtie2 version 2.3.4.3 (Langmead and Salzberg 2012) with default parameters. The resulting SAM files were converted to BAM files, sorted, and indexed using SAMtools version 1.9 (Danecek et al. 2021). We estimated the sequencing depth based on 10-kb sliding windows with 2-kb steps, and the sequencing depth of each window was estimated using Mosdepth version 0.2.3 (Pedersen and Quinlan 2018). We normalized the overall sequencing depths among male individuals and female individuals based on methods used in Li et al. 2020. The overall sequencing depth distribution was plotted using a violin plot in ggplot2 version 3.2.1 (Wickham 2016). The X chromosome assigned to the chromosome had about half the ratio of sequencing depth between males and females compared to the others.

### Comparative genomics and genome synteny analyses

We compared the genomes of the GWSS, *Nephotettix cincticeps* (rice green leafhopper) (Yan et al. 2021), and *Nilaparvata lugens* (brown planthopper) (Ye et al. 2021). The protein sequences of each genome were downloaded. We used OrthoVenn2 (Xu et al. 2019) with *e*-value = 1e-5 and inflation value = 1.5 to cluster orthologous groups and to create a Venn diagram of these three genomes. We used MCscanX (Wang et al. 2012) to evaluate the whole genome synteny between these species. All parameters were used as defaults in MCscanX. SynVisio (https://github.com/kiranbandi/synvisio) was used to display syntenies between genomes.

### Genomic distribution of brochosome-related genes

We used the list of brochosome-related genes as blastp queries to locate homologous sequences in the GWSS. We manually curated homologs and annotated them on our genome assembly using the ‘protein2genome’ mode of exonerate version 2.2.0 (Slater and Birney 2005). Similarly, we used reciprocal best blast hits of the GWSS brochosome-related genes to locate homologous sequences in the *Ne. cincticeps* genome. These homologous sequences were used as queries in the second round of blastp with the annotated proteins of *Ne. cincticeps* genome as the database. The location of each brochosome-related gene was shown in an ideogram produced by the R package Rideogram (Hao et al. 2020). The gene density was calculated with a sliding window size of 1Mb.

### RNAseq differential expression analyses of brochosome-related genes

To perform differential expression analyses of brochosome-related genes, we generated transcriptomes for Malpighian tubules and the rest of the organs of the abdomen of *H. vitripennis* with three biological replicates. For each paired set of transcriptomes, we used four frozen adult males for dissection and RNA extraction. The insects were dissected in a cold 1X PBS solution to remove all organs from the abdomen. All the Malpighian tubules were separated from the rest of the organs of the abdomen and pooled into a 1.7 mL microcentrifuge tube containing 200 μL PBS. The remaining organs were also pooled in 200 μL PBS. The RNA was then extracted from both samples using the Quick-RNA MiniPrep kit following the manufacturer’s instructions (Zymo Research, Irvine, CA). Total RNA in samples was quantified using a Qubit 2.0 Fluorometer (Life Technologies, Carlsbad, CA, USA), and RNA integrity was checked with a 4200 TapeStation (Agilent Technologies, Palo Alto, CA, USA). Sequencing libraries were quantified using a Qubit and validated using a TapeStation as well as by quantitative PCR (Applied Biosystems, Carlsbad, CA, USA). The sequencing libraries were multiplexed and clustered onto a flow cell and loaded on an Illumina HiSeq 4000 using a 2×150 paired-end (PE) configuration.

Differential gene expression analyses were performed to identify brochosome-related genes that are significantly upregulated in the Malpighian tubules. We used a list of candidate brochosome-related genes from a previous study that was based on proteomics and transcriptomics of the leafhopper *Graphocephala fennahi* (Rakitov et al. 2018). The protein sequences of these candidate genes were used as queries in blastp against all proteins inferred from our genome and from the other two GWSS genomes in GenBank. We used the same approach from the previous section for sequence read cleaning. The clean reads were mapped to the chromosome-level assembly using HISAT2 version 2.1.0 (Kim et al. 2019) with -k 3. We used featureCounts (Liao et al. 2014) to estimate the number of reads mapped to the exons of each candidate gene (“–type exon”). Counts of the genes were normalized, and we identified differentially expressed genes using DESeq2 version 1.20.0 in R (Love et al. 2014; R Core Team 2014). The Malpighian tubules were treated as one condition and the rest of the organs of the abdomen sample was used as the other condition. The Wald significance test was used to identify differentially expressed genes. Brochosome-related genes were identified as genes that were >2-fold upregulated in the Malpighian tubules compared to the other organs of the abdomen.

### Shotgun proteomics (LC-MS/MS) of Malpighian tubules

Multiple *H. vitripennis* individuals were collected from wild sunflowers (*Helianthus annuus*) near the entrance of the McKinney Roughs Nature Park at Cedar Creek, TX, and from crape myrtles (*Lagerstroemia indica*) near the campus of the University of Texas at Austin. To dissect the Malpighian tubules, female *H. vitripennis* individuals were frozen at -20°C for 5 min, then dissected in 1X PBS solution to extract the organs from the abdomen. Malpighian tubules were separated from the rest of the organs. The Malpighian tubules from four individuals were pooled into 1.7 mL microcentrifuge tubes containing 200 μL 1X PBS. For comparison, the remaining organs of the abdomen from two individuals (to achieve roughly equivalent protein concentration as the Malpighian tubules) were also pooled (Malpighian tubules: *n=4*; gut: *n=4*). All samples were then centrifuged at 4,000 x g for 10 min at 4°C to pellet the tissue. Following the removal of the supernatant, the pellet was homogenized with a pestle. Two different lysis buffers were added to the samples. For six samples (Malpighian tubules: *n=3*; gut: *n=3*), 300 μL of IP lysis buffer (25 mM Tris, 1 mM EDTA, 5% glycerol, 0.15 M NaCl, 1% Triton X-100) was added. For two samples (Malpighian tubules: *n=1*; gut: *n=1*), 300 μL of SDS lysis buffer (0.05% SDS, 5 mM EDTA pH 8.0, 1 mM PMSF, 10 mM DTT, 0.8 mg DNase I) was added. All samples were then centrifuged at 12,000 x g for 20 min at 4°C. The supernatants were transferred to clean microcentrifuge tubes and stored at -20°C until needed. Protein concentration was quantified with the Quick Start Bovine Serum Albumin Bradford assay according to the manufacturer’s protocol (Bio-Rad Laboratories, Inc., Hercules, CA) and using an Eppendorf Biophotometer 6131.

In preparation for protein electrophoresis, Malpighian tubule samples were diluted in water to attain appropriate protein concentrations, giving a total of 15 μL volume per sample. 3 μL 6X SDS loading dye containing reducing agent (0.6 M DTT,0.35 M Tris pH 6.8, 30% v/v glycerol, 10% w/v SDS, 0.012% w/v bromophenol blue) was added to each sample. Samples were then heated at 100°C for 10 min, allowed to cool to room temperature, and centrifuged for 30 s. The samples were run in 12% polyacrylamide gel for 20 min at 70 V and a constant 30 mA. The gel was then stained with Coomassie G-250 for 30 min and destained with 10% ethanol and 5% acetic acid solution overnight. Sample lanes were cut from the gel and stored in the same destain solution at 4°C until trypsin digestion.

The LC-MS/MS shotgun proteomics was performed by the Proteomics Core at UT Austin. Raw LC-MS/MS spectra were processed using Proteome Discoverer (v2.3) (Orsburn 2021). We used the Percolator node in Proteome Discoverer to assign unique peptide spectral matches (PSMs) at FDR <5% to the composite form of the GWSS reference proteome which comprises 31,235 proteins. In order to identify proteins statistically significantly associated with each bait, we calculated both a log2 fold-change and a Z-score for each protein based on the observed PSMs in the bait (‘expt’) versus control (‘ctrl’) pulldown. The fold-change was computed for each protein according to the methods reported in a recent study (Lee et al. 2020).

### Shotgun proteomics (LC-MS/MS) of purified brochosomes

To obtain purified brochosomes of *H. vitripennis*, five female individuals were stored at -80°C and soaked in acetone for 25 min. The solution was then sonicated using a Bransonic 2800 bath sonicator for 3 min to suspend the brochosome particles in the solution. The brochosome solution was then filtered using a glass fiber syringe filter with a pore size of 1 μm to remove impurities and leafhopper fragments. To further concentrate the brochosome solution, the filtered solution was centrifuged using an Eppendorf 5804R centrifuge at 16,000 x g for 25 min. After centrifugation, the supernatant was removed and the concentrated product was stored in ambient conditions.

In preparation for protein electrophoresis, the dried pellet samples were resuspended in 18 μL of Laemmli’s buffer (50 mM Tris-HCl pH 8.8, 2% SDS, 10% glycerol, 0.01% bromophenol blue, 100 mM DTT) (Rakitov et al. 2018). They were then heated at 100°C for 10 min, allowed to cool to room temperature, and centrifuged for 30s. The samples were then run on a protein gel and further analyzed in the same manner as the Malpighian tubule samples.

### Identifying symbiosis-related genes with differential expression analyses

Using transcriptome data from a previous study (Mao and Bennett 2020), we compared transcriptomes of three tissue types: red bacteriome, yellow bacteriome, and the remainder of the body, with three biological replicates for each tissue type. These data were downloaded through the AWS links on Genbank (BioProject: PRJNA342859, Accession: SRR10060917, SRR10060918, SRR10060919). We used Trimmomatic version 0.38 (Bolger et al. 2014) to remove low-quality reads. Filtered reads were mapped to our chromosome-level assembly using HISAT2 version 2.1.0 (Kim et al.2019) with -k 3. We used featureCounts (Liao et al. 2014) to estimate the number of reads mapped to the exons of each gene (“–type exon”). Counts of the genes were normalized, and we identified differentially expressed genes using DESeq2 version 1.20.0 in R (Love et al. 2014; R Core Team 2014). To take into account biases due to variation in sequencing depth and RNA composition, the median of ratios method implemented in DESeq2 was used for normalization. Given that there were three tissue types, two differential analyses were performed. Each bacteriome type was treated as one condition and the body as the other condition. The Wald significance test was used to identify differentially expressed genes. Symbiosis-related genes were identified as genes that were >2-fold upregulated in the two bacteriomes compared to the body. The statistical significance of overlapping upregulated host genes between two bacteriomes was tested with a hypergeometric test in package ‘stats’ version 3.6.1 in R (R Core Team 2014).

### Evolutionary history of HGT genes

Previous studies identified multiple bacterial genes that were horizontally transferred from bacteria into leafhoppers using the draft genomes of *H. vitripennis* and *M. quadrilineatus* (Mao et al. 2018; Mao and Bennett 2020). To understand the evolutionary sources of HGT genes previously identified in these two leafhoppers, we used their translated sequences of HGT genes as blastp queries. We also assembled the translated protein sequences of 25 hemipteran genomes and 102 transcriptomes as our blast database. A blastp search with an E-value of 1e -5, a bit-score of 100, and percent identity of 40% as thresholds were used to find homologous protein sequences of queries from the blast database. All protein sequences from the blastp hits were clustered into gene families using OrthoFinder version 2.3.12 (Emms and Kelly 2019) with defaults. The bacterial protein sequences used as outgroups in previous studies (Mao et al. 2018; Mao and Bennett 2020) were also downloaded from Genbank. For each HGT gene family plus outgroup sequences, homologous protein sequences were aligned using MUSCLE v3.8.31 (Edgar 2004) with default parameters. Sequences with low overlap in the protein alignments were removed by using -resoverlap 0.75-seqoverlap 60 in trimAl v1.2 (Capella-Gutiérrez et al. 2009). The ‘-gappyout’ option was used to remove columns with many gaps from the protein sequence alignments. We then manually inspected each protein alignment and removed any ambiguous sequences. Finally, gene trees were built with IQ-TREE multicore version 1.6.1 with 1000 bootstrap replicates using models selected by MFP ModelFinder (Nguyen et al. 2015).

## Data Availability

*Homalodisca vitripennis* genome assembly and sequencing data are available on NCBI under the BioProject ID PRJNA776266. RefSeq genome annotation is available at: https://ftp.ncbi.nlm.nih.gov/genomes/all/GCF/021/130/785/GCF_021130785.1_UT_GW SS_2.1/; Annotation information is available at: https://www.ncbi.nlm.nih.gov/genome/annotation_euk/Homalodisca_vitripennis/100/; Genome information is available at: https://www.ncbi.nlm.nih.gov/genome/?term=txid197043[orgn]

## Acknowledgments

This work was supported by the US Army Research Office MURI award W911NF-20-1-0195. Zheng Li is supported by the National Science Foundation Postdoctoral Research Fellowships in Biology Program under grant DBI-2109306. We thank Tom Smith, Laila Phillips, and Eli Powell from the Moran lab for assisting with insect dissection, and RNA and protein extractions. We thank Sarah Bialik from the Barrick lab for the SEM imaging and Mostafa Nasser from the Freemen lab at UT Austin for purifying brochosomes. We thank Maria Person, Michelle Gadush, and Peter Faull at the Proteomics Facility at UT Austin. We also thank Gordon Bennett and Meng Mao for assisting in accessing their data.

