## Supplemental Figure S1-6 for "The genomic basis of evolutionary novelties in a leafhopper"

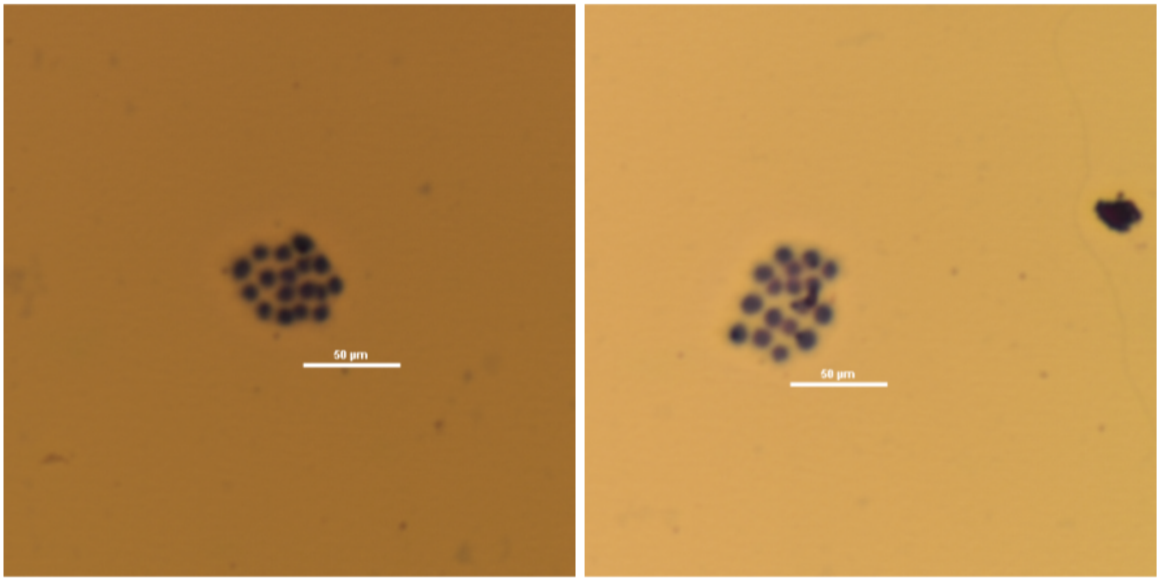


**Fig S1.** Conventional staining with Giemsa of metaphase I cells for chromosomes of a male GWSS.

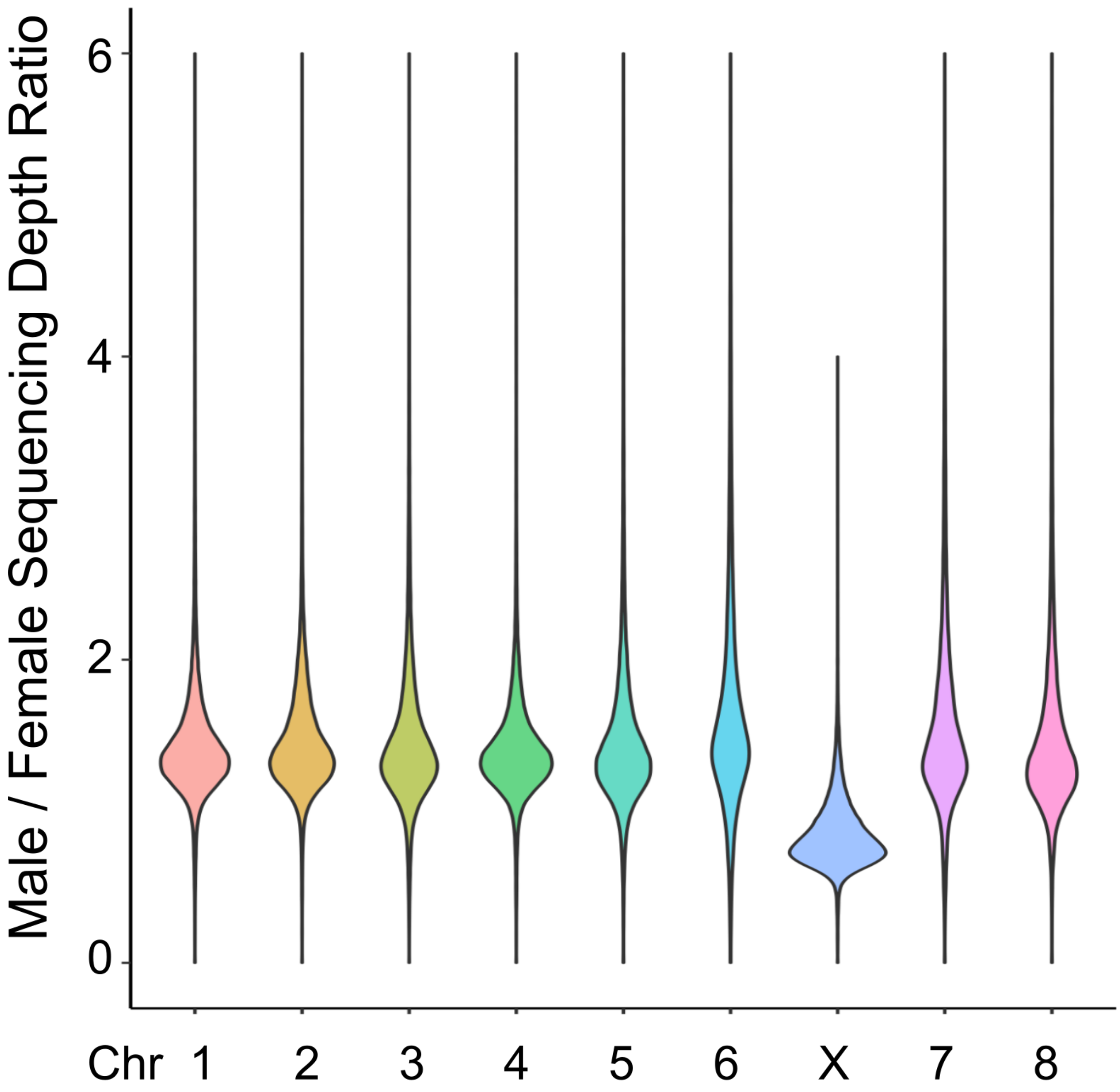


**Fig S2.** Normalized sequencing depth between male and females of 8 autosomes and the X chromosome. The X chromosome showed about half of the sequencing depth ratio between sexes compared to the ratios found for autosomes.

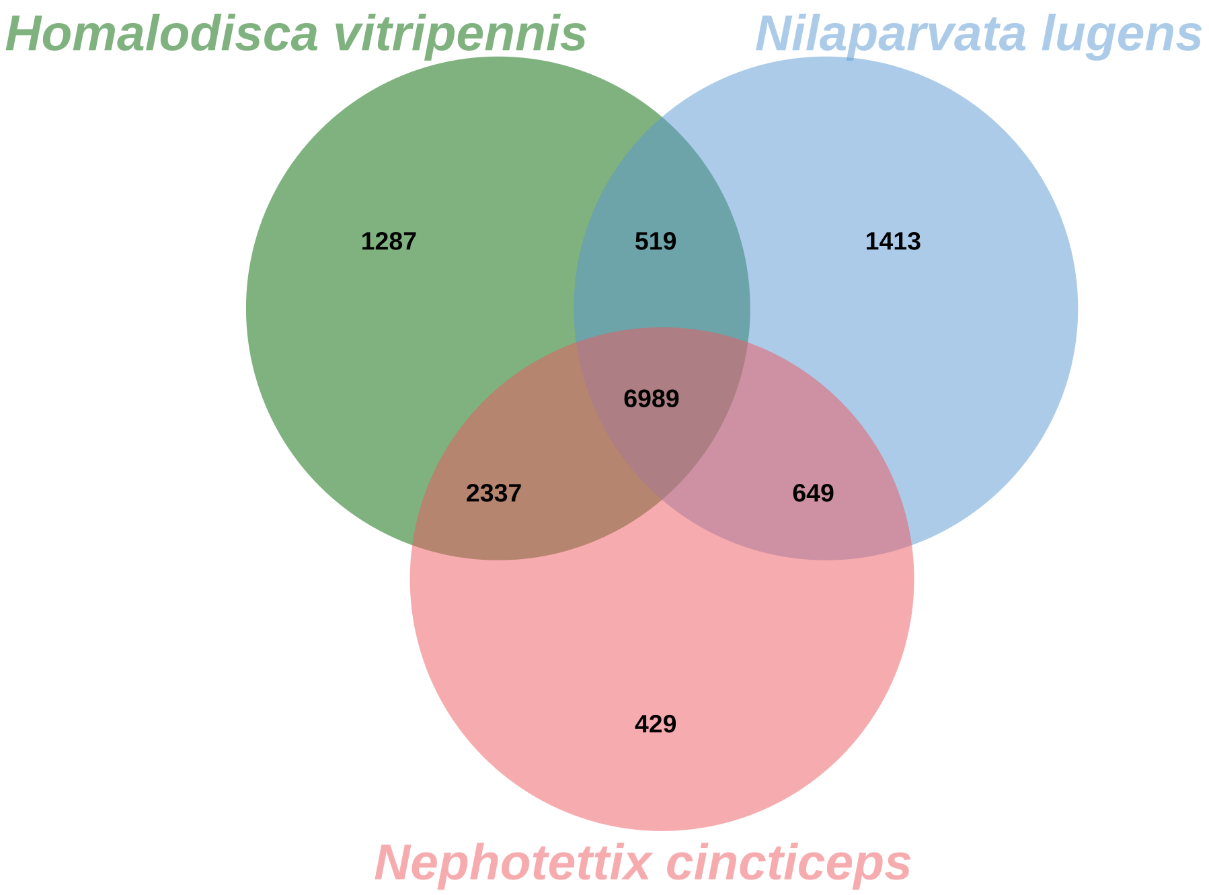


**Fig S3.** The Venn diagram of orthologous clusters between *Homalodisca vitripennis* (GWSS), *Nephotettix cincticeps* (rice green leafhopper), and *Nilaparvata lugens* (brown planthopper).

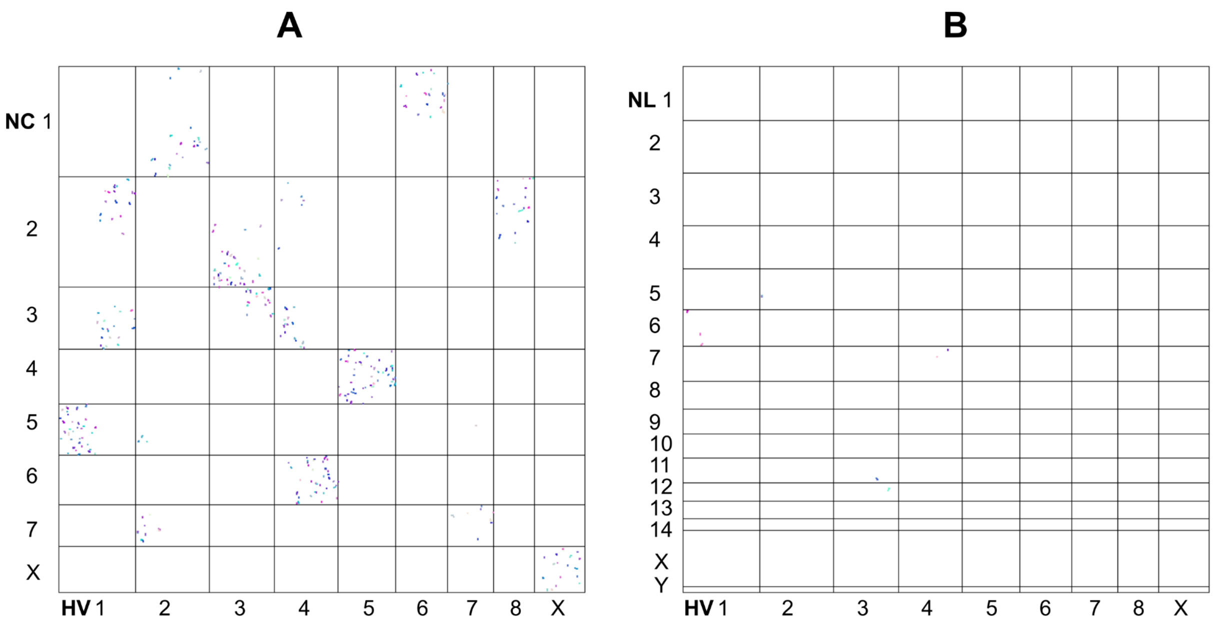


**Fig S4.** Genome dot plots of three available chromosome-level genome assemblies of leafhoppers and planthoppers. Different colors of dots represent different syntenic blocks. **A:** *Homalodisca vitripennis* vs *Nephotettix cincticeps* (rice green leafhopper) and **B:** *Homalodisca vitripennis* vs. *Nilaparvata lugens* (brown planthopper).

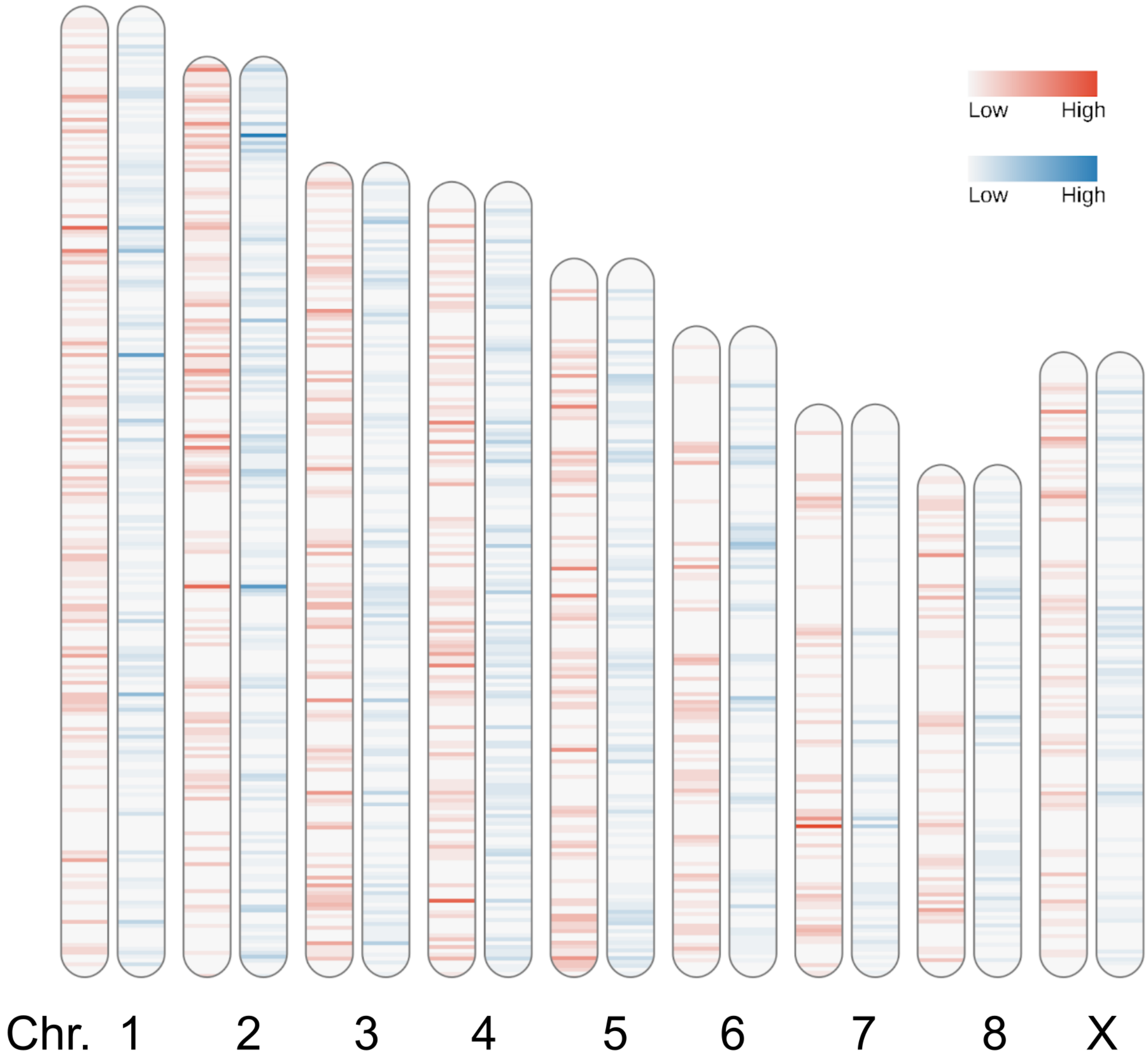


**Fig S5.** The chromosomal location and gene density of symbiosis-related genes in the GWSS. The intensity of the color on each chromosome represents gene density. The red color corresponds to genes upregulated in the more ancestral yellow bacteriome. The blue corresponds to genes upregulated in the more recently evolved red bacteriome.

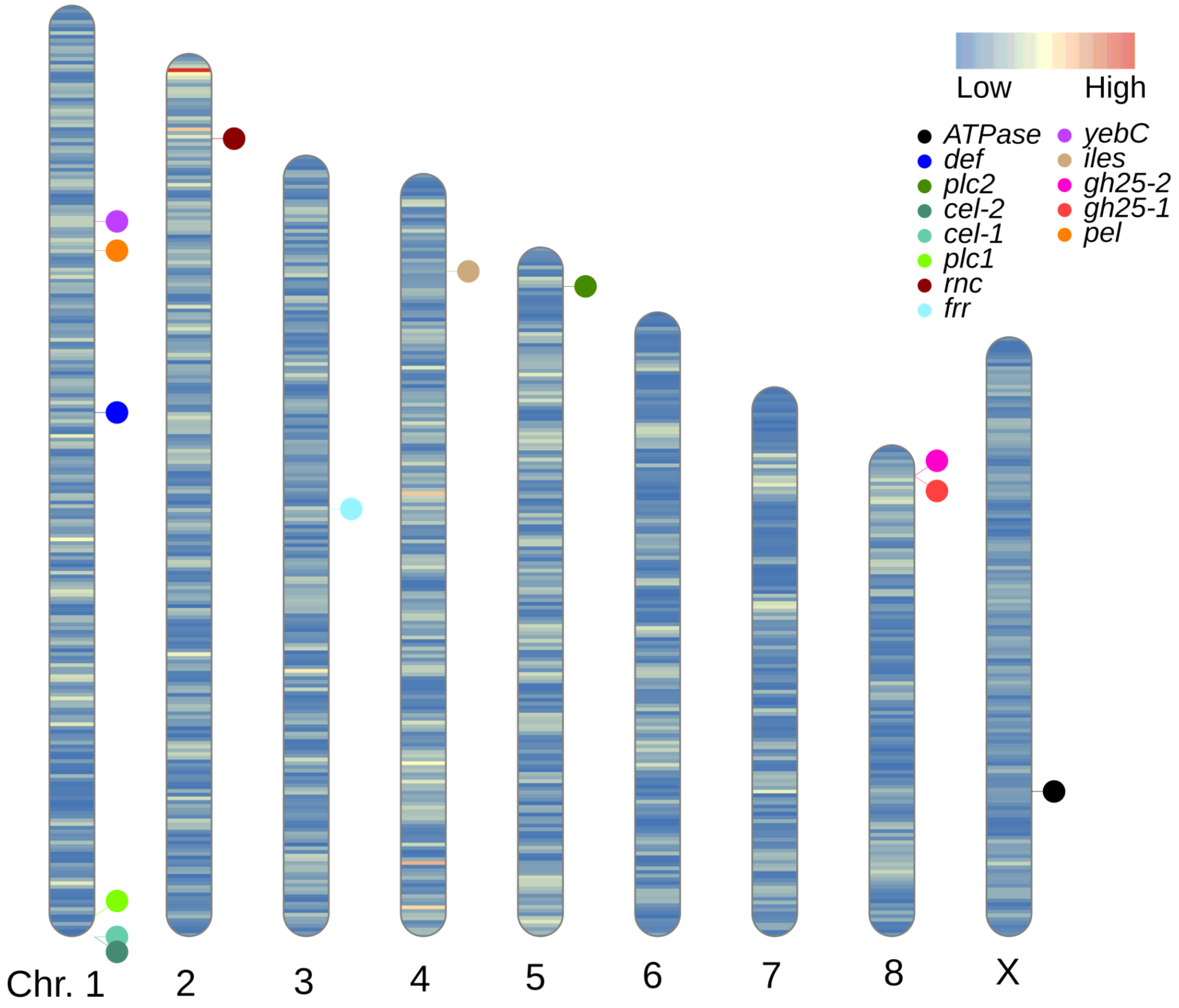


**Fig S6.** Ideogram with chromosomal locations of HGT genes in the GWSS genomes. The intensity of the color on each chromosome represents gene density.

**Fig S7.** Phylogenies for 16 horizontally transferred genes in Auchenorrhyncha. The bootstrap values are shown on the node. Sequences of Auchenorrhyncha are shown in black; Sequences of other Hemipteran insects are shown in red; The bacterial outgroup sequences are shown in blue.
