## Supplemental Figure S7 for "The genomic basis of evolutionary novelties in a leafhopper"

*alv*

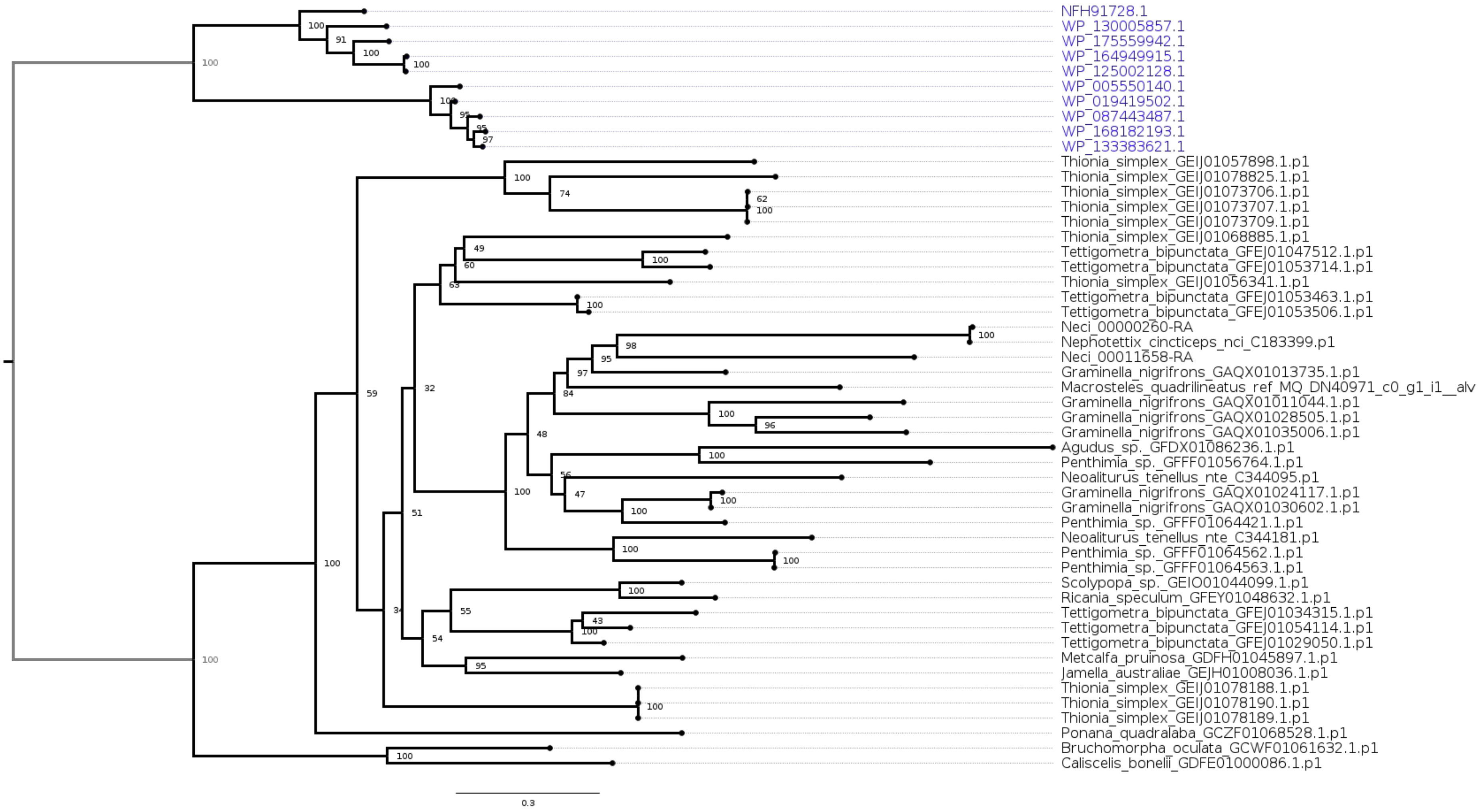
NFH91728.1
WP_130005857.1
WP_175559942.1
WP_164949915.1
WP_125002128.1
WP_005550140.1
WP_019419502.1
WP_087443487.1
WP_168182193.1
WP_133383621.1
Thionia_simplex_GEIJ01057898.1.p1
Thionia_simplex_GEIJ01078825.1.p1
Thionia_simplex_GEIJ01073706.1.p1
Thionia_simplex_GEIJ01073707.1.p1
Thionia_simplex_GEIJ01073709.1.p1
Thionia_simplex_GEIJ01068885.1.p1
Tettigometra_bipunctata_GFEJ01047512.1.p1
Tettigometra_bipunctata_GFEJ01053714.1.p1
Thionia_simplex_GEIJ01056341.1.p1
Tettigometra_bipunctata_GFEJ01053463.1.p1
Tettigometra_bipunctata_GFEJ01053506.1.p1
Neci_00000260-RA
Nephotettix_cincticeps_nci_C183399.p1
Neci_00011658-RA
Graminella_nigrifrons_GAQX01013735.1.p1
Macrosteles_quadrilineatus_ref_MQ_DN40971_c0_g1_i1__alv
Graminella_nigrifrons_GAQX01011044.1.p1
Graminella_nigrifrons_GAQX01028505.1.p1
Graminella_nigrifrons_GAQX01035006.1.p1
Agudus_sp._GFDX01086236.1.p1
Penthimia_sp._GFFF01056764.1.p1
Neoaliturus_tenellus_nte_C344095.p1
Graminella_nigrifrons_GAQX01024117.1.p1
Graminella_nigrifrons_GAQX01030602.1.p1
Penthimia_sp._GFFF01064421.1.p1
Neoaliturus_tenellus_nte_C344181.p1
Penthimia_sp._GFFF01064562.1.p1
Penthimia_sp._GFFF01064563.1.p1
Scolypopa_sp._GEIO01044099.1.p1
Ricania_speculum_GFEY01048632.1.p1
Tettigometra_bipunctata_GFEJ01034315.1.p1
Tettigometra_bipunctata_GFEJ01054114.1.p1
Tettigometra_bipunctata_GFEJ01029050.1.p1
Metcalfa_pruinosa_GDFH01045897.1.p1
Jamella_australiae_GEJH01008036.1.p1
Thionia_simplex_GEIJ01078188.1.p1
Thionia_simplex_GEIJ01078190.1.p1
Thionia_simplex_GEIJ01078189.1.p1
Ponana_quadralaba_GCZF01068528.1.p1
Bruchomorpha_oculata_GCWF01061632.1.p1
Caliscelis_bonellii_GDFE01000086.1.p1
0.3

### *ATPase*

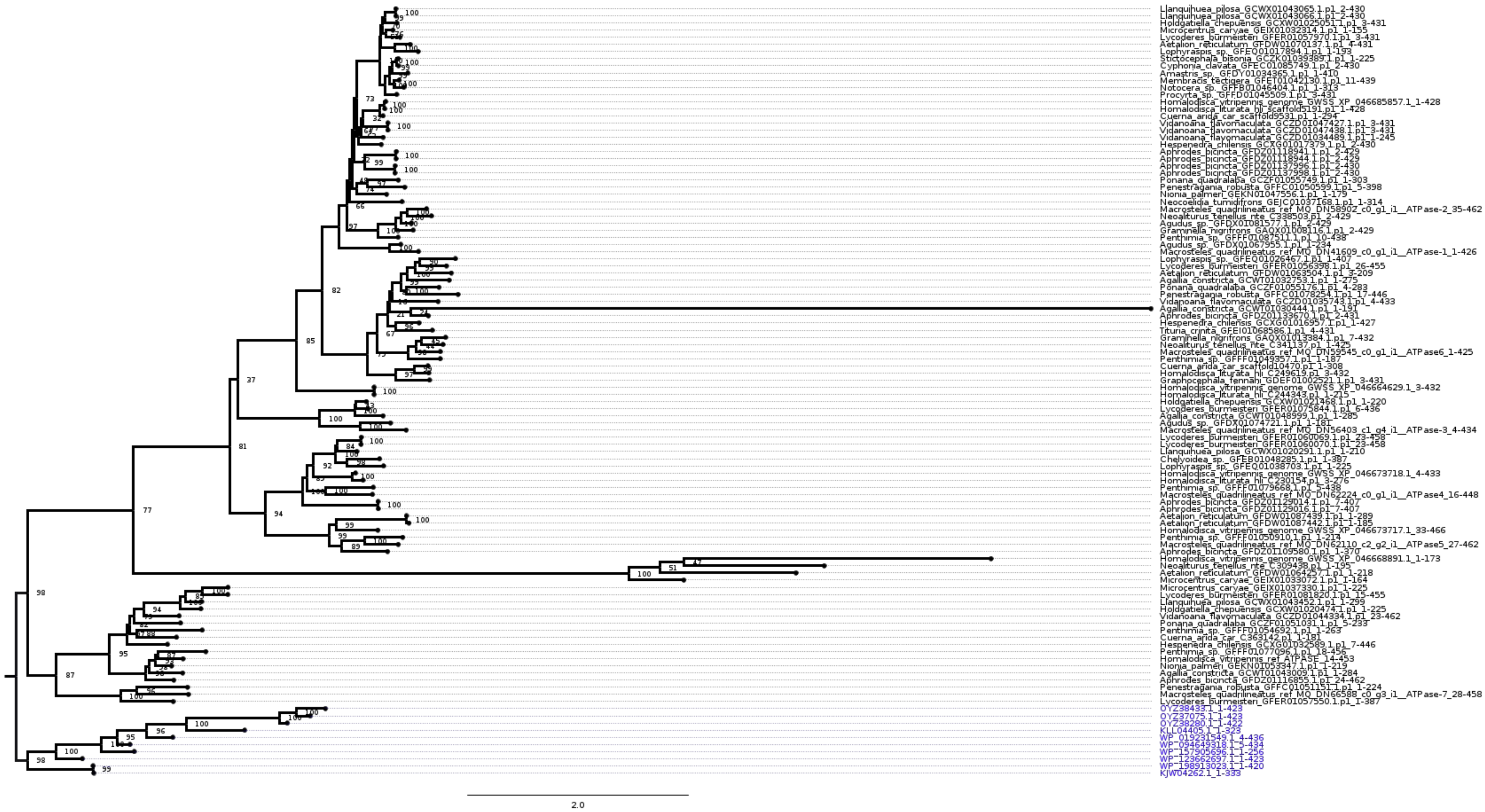
Llanquihuea_pilosa_GCWX01043065.1.p1_2-430
Llanquihuea_pilosa_GCWX01043066.1.p1_2-430
Holdgatiella_chepuensis_GCXW01025051.1.p1_3-431
Microcentrus_caryae_GEIX01032314.1.p1_1-155
Lycoderes_burmeisteri_GFER01057970.1.p1_3-431
Aetalion_reticulatum_GFDW01070137.1.p1_4-431
Lophyraspis_sp._GFEQ01017894.1.p1_1-193
Stictocephala_bisonia_GCZK01039389.1.p1_1-225
Cyphonia_clavata_GFEC01085749.1.p1_2-430
Amastris_sp._GFDY01034365.1.p1_1-410
Membracis_tectigera_GFET01042130.1.p1_11-439
Notocera_sp._GFFB01046404.1.p1_1-313
Procyrta_sp._GFFD01045509.1.p1_3-431
Homalodisca_vitripennis_genome_GWSS_XP_046685857.1_1-428
Homalodisca_liturata_hli_scaffold5191.p1_1-428
Cuerna_arida_car_scaffold9531.p1_1-294
Vidanoana_flavomaculata_GCZD01047427.1.p1_3-431
Vidanoana_flavomaculata_GCZD01047438.1.p1_3-431
Vidanoana_flavomaculata_GCZD01034489.1.p1_1-245
Hespenedra_chilensis_GCXG01017379.1.p1_2-430
Aphrodes_bicincta_GFDZ01118941.1.p1_2-429
Aphrodes_bicincta_GFDZ01118944.1.p1_2-429
Aphrodes_bicincta_GFDZ01137996.1.p1_2-430
Aphrodes_bicincta_GFDZ01137998.1.p1_2-430
Ponana_quadralaba_GCZF01055749.1.p1_1-303
Penestragania_robusta_GFFC01050599.1.p1_5-398
Nionia_palmeri_GEKN01047556.1.p1_1-179
Neocoelidia_tumidifrons_GEJC01037168.1.p1_1-314
Macrosteles_quadrilineatus_ref_MQ_DN58902_c0_g1_i1__ATPase-2_35-462
Neoaliturus_tenellus_nte_C338503.p1_2-429
Agudus_sp._GFDX01081577.1.p1_2-429
Graminella_nigrifrons_GAQX01008116.1.p1_2-429
Penthimia_sp._GFFF01087511.1.p1_10-438
Agudus_sp._GFDX01067955.1.p1_1-234
Macrosteles_quadrilineatus_ref_MQ_DN41609_c0_g1_i1__ATPase-1_1-426
Lophyraspis_sp._GFEQ01026467.1.p1_1-407
Lycoderes_burmeisteri_GFER01056398.1.p1_26-455
Aetalion_reticulatum_GFDW01063504.1.p1_3-209
Agallia_constricta_GCWT01032753.1.p1_1-275
Ponana_quadralaba_GCZF01055176.1.p1_4-283
Penestragania_robusta_GFFC01078254.1.p1_17-446
Vidanoana_flavomaculata_GCZD01035743.1.p1_4-433
Agallia_constricta_GCWT01030444.1.p1_1-191
Aphrodes_bicincta_GFDZ01133670.1.p1_2-431
Hespenedra_chilensis_GCXG01016957.1.p1_1-427
Tituria_crinita_GFEI01068586.1.p1_4-431
Graminella_nigrifrons_GAQX01013384.1.p1_7-432
Neoaliturus_tenellus_nte_C341137.p1_1-425
Macrosteles_quadrilineatus_ref_MQ_DN59545_c0_g1_i1__ATPase6_1-425
Penthimia_sp._GFFF01049357.1.p1_1-187
Cuerna_arida_car_scaffold10470.p1_1-308
Homalodisca_liturata_hli_C249619.p1_3-432
Graphocephala_fennahi_GDEF01002521.1.p1_3-431
Homalodisca_vitripennis_genome_GWSS_XP_046664629.1_3-432
Homalodisca_liturata_hli_C244343.p1_1-215
Holdgatiella_chepuensis_GCXW01021468.1.p1_1-220
Lycoderes_burmeisteri_GFER01075844.1.p1_6-436
Agallia_constricta_GCWT01048999.1.p1_1-285
Agudus_sp._GFDX01074721.1.p1_1-181
Macrosteles_quadrilineatus_ref_MQ_DN56403_c1_g4_i1__ATPase-3_4-434
Lycoderes_burmeisteri_GFER01060069.1.p1_23-458
Lycoderes_burmeisteri_GFER01060070.1.p1_23-458
Llanquihuea_pilosa_GCWX01020291.1.p1_1-210
Chelyoidea_sp._GFEB01048285.1.p1_1-387
Lophyraspis_sp._GFEQ01038703.1.p1_1-225
Homalodisca_vitripennis_genome_GWSS_XP_046673718.1_4-433
Homalodisca_liturata_hli_C230154.p1_3-276
Penthimia_sp._GFFF01079668.1.p1_5-438
Macrosteles_quadrilineatus_ref_MQ_DN62224_c0_g1_i1__ATPase4_16-448
Aphrodes_bicincta_GFDZ01129014.1.p1_7-407
Aphrodes_bicincta_GFDZ01129016.1.p1_7-407
Aetalion_reticulatum_GFDW01087439.1.p1_1-289
Aetalion_reticulatum_GFDW01087442.1.p1_1-185
Homalodisca_vitripennis_genome_GWSS_XP_046673717.1_33-466
Penthimia_sp._GFFF01050910.1.p1_1-214
Macrosteles_quadrilineatus_ref_MQ_DN62110_c2_g2_i1__ATPase5_27-462
Aphrodes_bicincta_GFDZ01109580.1.p1_1-370
Homalodisca_vitripennis_genome_GWSS_XP_046668891.1_1-173
Neoaliturus_tenellus_nte_C309438.p1_1-195
Aetalion_reticulatum_GFDW01064257.1.p1_1-218
Microcentrus_caryae_GEIX01033072.1.p1_1-164
Microcentrus_caryae_GEIX01037330.1.p1_1-225
Lycoderes_burmeisteri_GFER01081820.1.p1_15-455
Llanquihuea_pilosa_GCWX01043452.1.p1_1-299
Holdgatiella_chepuensis_GCXW01020474.1.p1_1-225
Vidanoana_flavomaculata_GCZD01044334.1.p1_23-462
Ponana_quadralaba_GCZF01051031.1.p1_5-233
Penthimia_sp._GFFF01054692.1.p1_1-263
Cuerna_arida_car_C363142.p1_1-181
Hespenedra_chilensis_GCXG01032589.1.p1_7-446
Penthimia_sp._GFFF01077096.1.p1_18-456
Homalodisca_vitripennis_ref_ATPASE_14-453
Nionia_palmeri_GEKN01053347.1.p1_1-219
Agallia_constricta_GCWT01043009.1.p1_1-284
Aphrodes_bicincta_GFDZ01116855.1.p1_24-462
Penestragania_robusta_GFFC01051151.1.p1_1-224
Macrosteles_quadrilineatus_ref_MQ_DN66588_c0_g3_i1__ATPase-7_28-458
Lycoderes_burmeisteri_GFER01057550.1.p1_1-387
OYZ38433.1_1-423
OYZ37075.1_1-423
OYZ38280.1_1-422
KLL04405.1_1-323
WP_019231549.1_4-436
WP_094649318.1_5-434
WP_157905696.1_1-256
WP_123662697.1_1-423
WP_198913023.1_1-420
KJW04262.1_1-333
2.0

*cel*

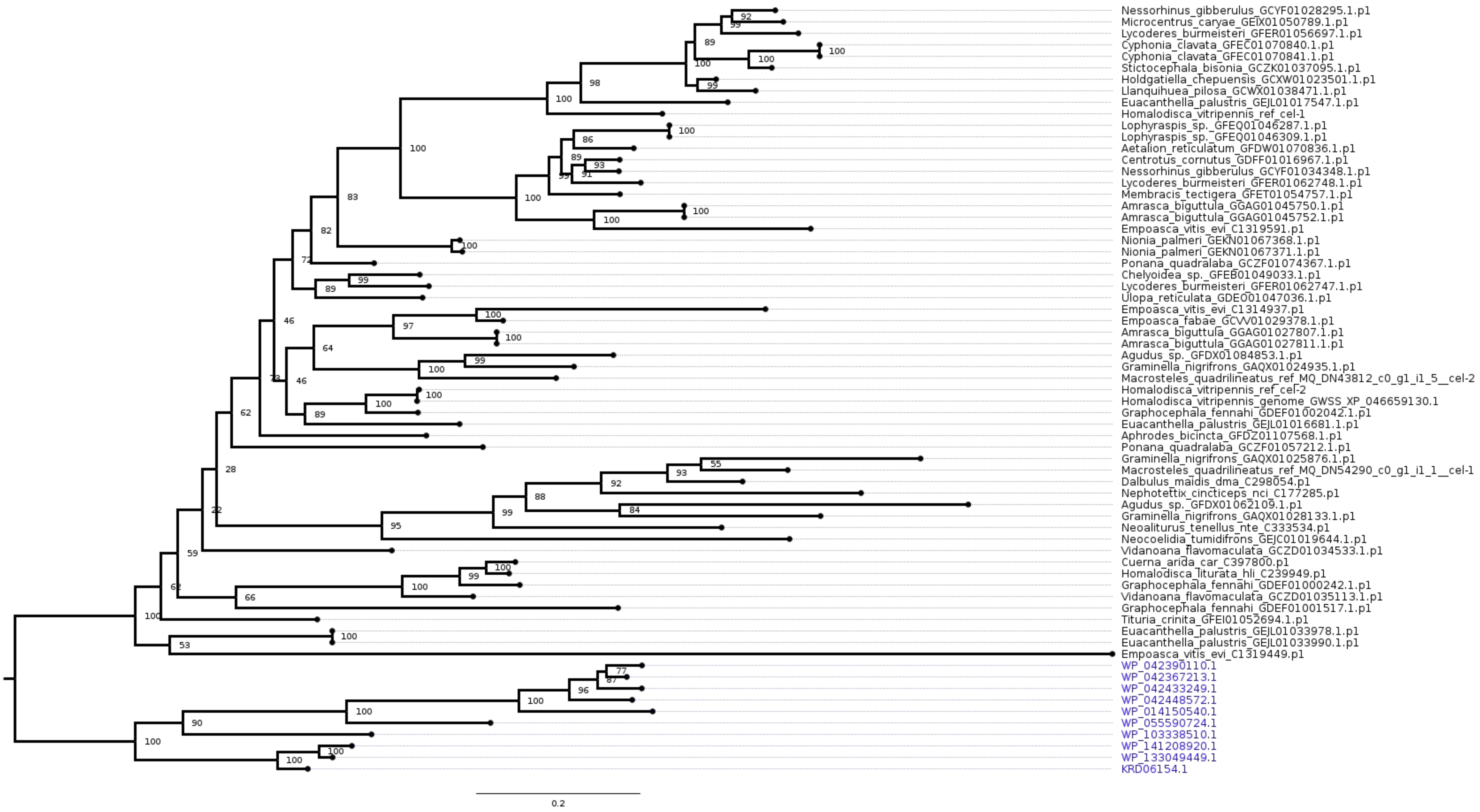
Nessorhinus_gibberulus_GCYF01028295.1.p1
Microcentrus_caryae_GEIX01050789.1.p1
Lycoderes_burmeisteri_GFER01056697.1.p1
Cyphonia_clavata_GFEC01070840.1.p1
Cyphonia_clavata_GFEC01070841.1.p1
Stictocephala_bisonia_GCZK01037095.1.p1
Holdgatiella_chepuensis_GCXW01023501.1.p1
Llanquihuea_pilosa_GCWX01038471.1.p1
Euacanthella_palustris_GEJL01017547.1.p1
Homalodisca_vitripennis_ref_cel-1
Lophyraspis_sp._GFEQ01046287.1.p1
Lophyraspis_sp._GFEQ01046309.1.p1
Aetalion_reticulatum_GFDW01070836.1.p1
Centrotus_cornutus_GDFF01016967.1.p1
Nessorhinus_gibberulus_GCYF01034348.1.p1
Lycoderes_burmeisteri_GFER01062748.1.p1
Membracis_tectigera_GFET01054757.1.p1
Amrasca_biguttula_GGAG01045750.1.p1
Amrasca_biguttula_GGAG01045752.1.p1
Empoasca_vitis_evi_C1319591.p1
Nionia_palmeri_GEKN01067368.1.p1
Nionia_palmeri_GEKN01067371.1.p1
Ponana_quadralaba_GCZF01074367.1.p1
Chelyoidea_sp._GFEB01049033.1.p1
Lycoderes_burmeisteri_GFER01062747.1.p1
Ulopa_reticulata_GDEO01047036.1.p1
Empoasca_vitis_evi_C1314937.p1
Empoasca_fabae_GCVV01029378.1.p1
Amrasca_biguttula_GGAG01027807.1.p1
Amrasca_biguttula_GGAG01027811.1.p1
Agudus_sp._GFDX01084853.1.p1
Graminella_nigrifrons_GAQX01024935.1.p1
Macrosteles_quadrilineatus_ref_MQ_DN43812_c0_g1_i1_5__cel-2
Homalodisca_vitripennis_ref_cel-2
Homalodisca_vitripennis_genome_GWSS_XP_046659130.1
Graphocephala_fennahi_GDEF01002042.1.p1
Euacanthella_palustris_GEJL01016681.1.p1
Aphrodes_bicincta_GFDZ01107568.1.p1
Ponana_quadralaba_GCZF01057212.1.p1
Graminella_nigrifrons_GAQX01025876.1.p1
Macrosteles_quadrilineatus_ref_MQ_DN54290_c0_g1_i1_1__cel-1
Dalbulus_maidis_dma_C298054.p1
Nephotettix_cincticeps_nci_C177285.p1
Agudus_sp._GFDX01062109.1.p1
Graminella_nigrifrons_GAQX01028133.1.p1
Neoaliturus_tenellus_nte_C333534.p1
Neocoelidia_tumidifrons_GEJC01019644.1.p1
Vidanoana_flavomaculata_GCZD01034533.1.p1
Cuerna_arida_car_C397800.p1
Homalodisca_liturata_hli_C239949.p1
Graphocephala_fennahi_GDEF01000242.1.p1
Vidanoana_flavomaculata_GCZD01035113.1.p1
Graphocephala_fennahi_GDEF01001517.1.p1
Tituria_crinita_GFEI01052694.1.p1
Euacanthella_palustris_GEJL01033978.1.p1
Euacanthella_palustris_GEJL01033990.1.p1
Empoasca_vitis_evi_C1319449.p1
WP_042390110.1
WP_042367213.1
WP_042433249.1
WP_042448572.1
WP_014150540.1
WP_055590724.1
WP_103338510.1
WP_141208920.1
WP_133049449.1
KRD06154.1
0.2

*def*

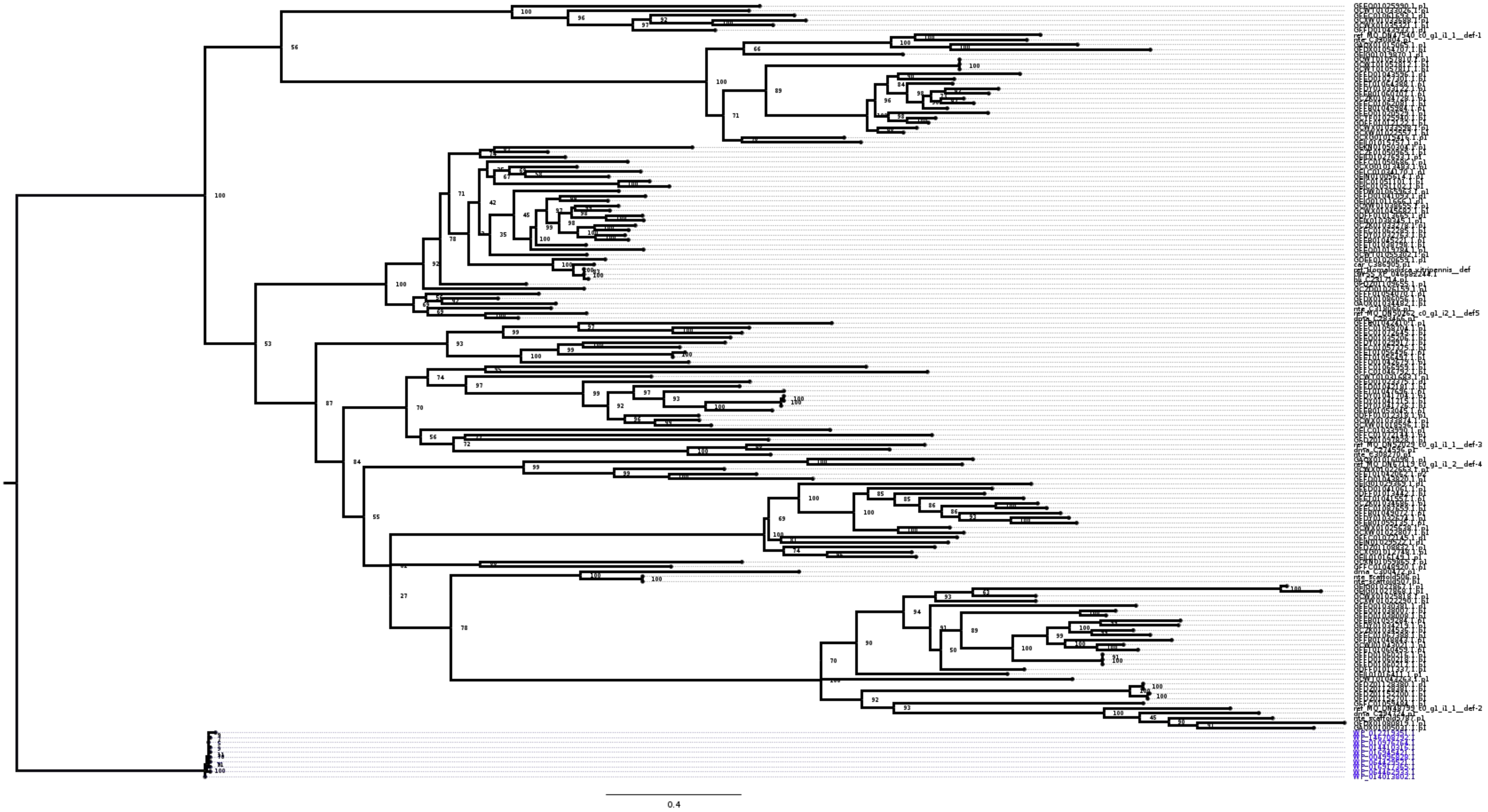

### *dut*

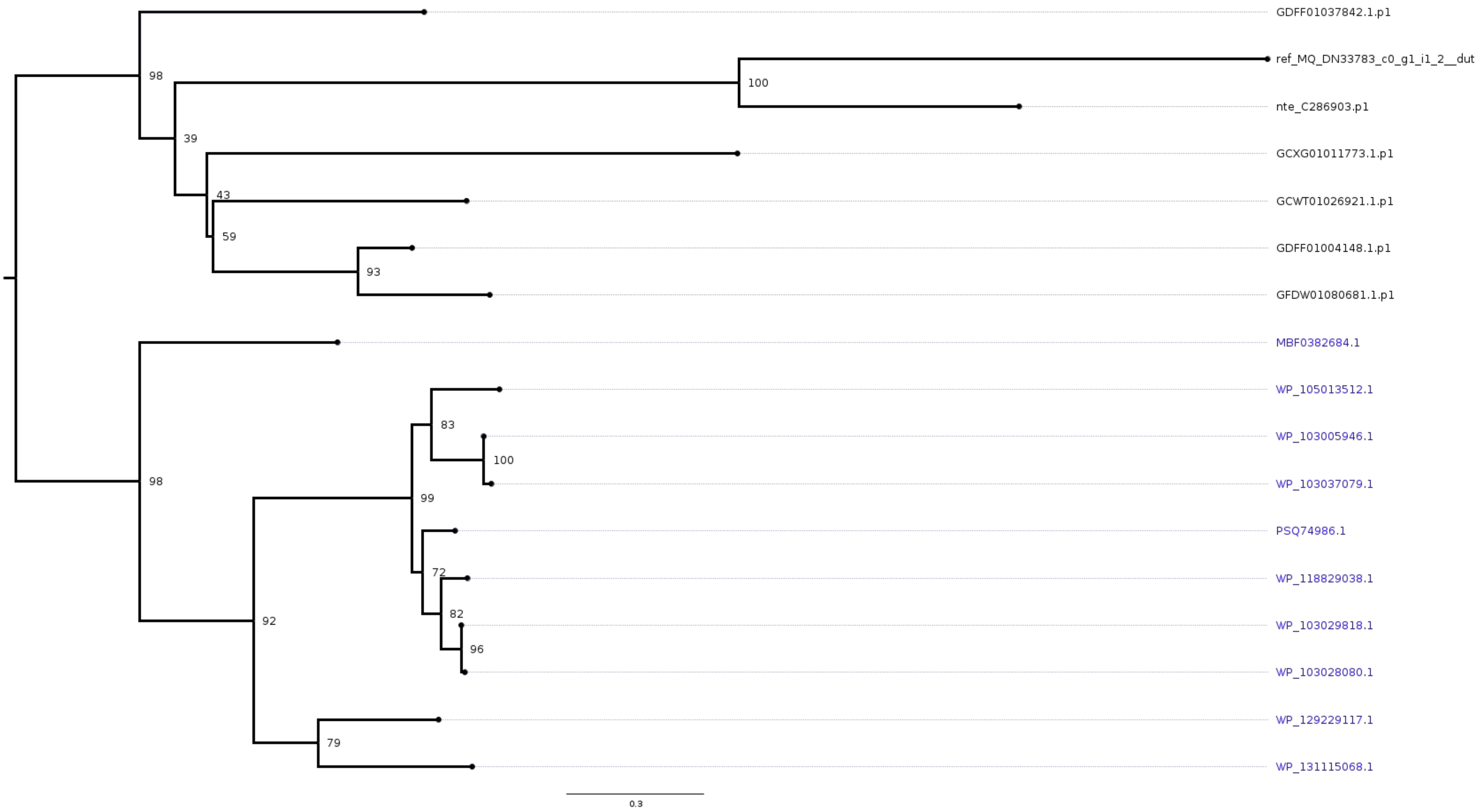
GDFF01037842.1.p1
ref_MQ_DN33783_c0_g1_i1_2__dut
nte_C286903.p1
GCXG01011773.1.p1
GCWT01026921.1.p1
GDFF01004148.1.p1
GFDW01080681.1.p1
MBF0382684.1
WP_105013512.1
WP_103005946.1
WP_103037079.1
PSQ74986.1
WP_118829038.1
WP_103029818.1
WP_103028080.1
WP_129229117.1
WP_131115068.1
98
100
39
43
59
93
98
83
100
99
72
82
96
92
79
0.3

### *frr*

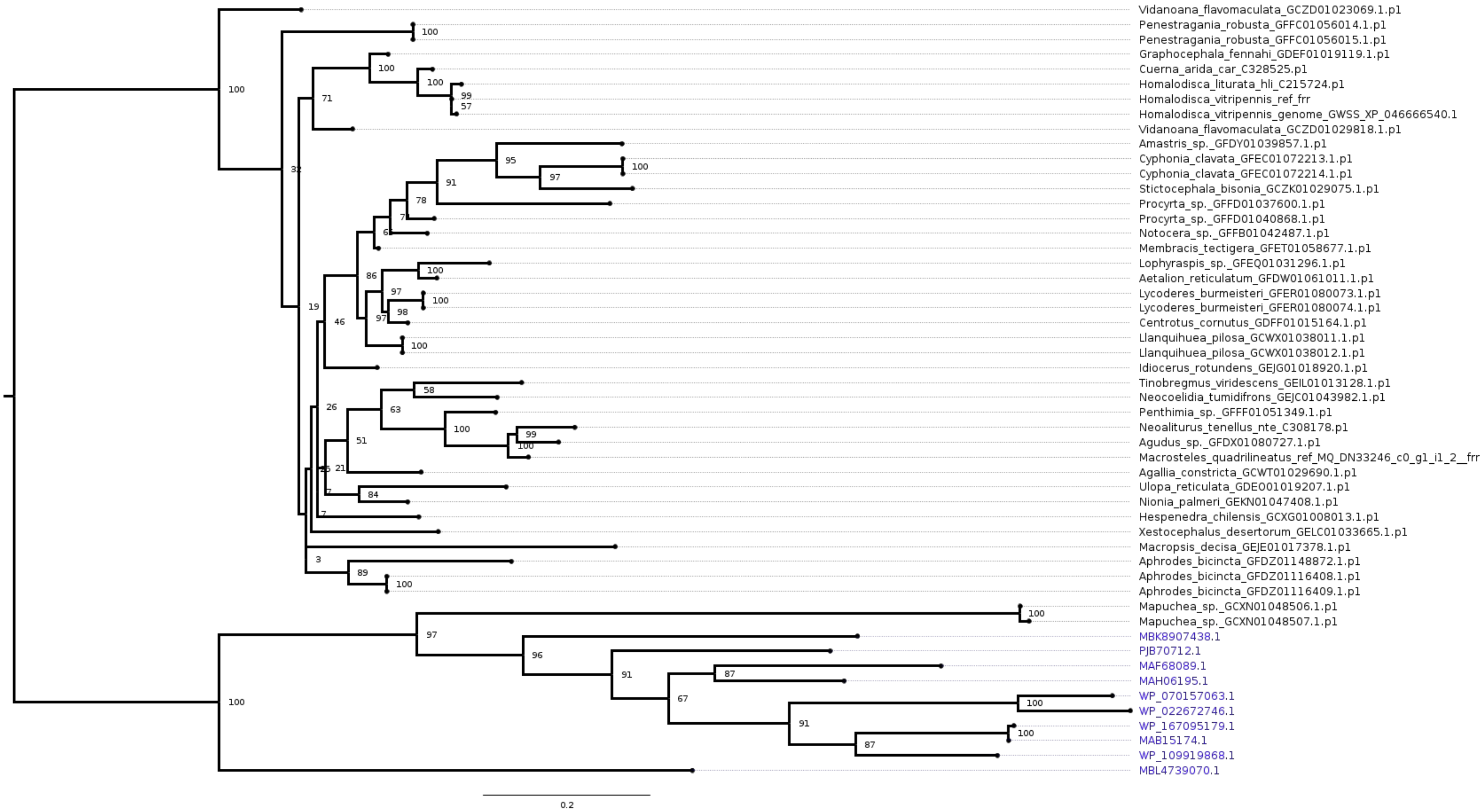
Vidanoana_flavomaculata_GCZD01023069.1.p1
Penestragania_robusta_GFFC01056014.1.p1
Penestragania_robusta_GFFC01056015.1.p1
Graphocephala_fennahi_GDEF01019119.1.p1
Cuerna_arida_car_C328525.p1
Homalodisca_liturata_hli_C215724.p1
Homalodisca_vitripennis_ref_frr
Homalodisca_vitripennis_genome_GWSS_XP_046666540.1
Vidanoana_flavomaculata_GCZD01029818.1.p1
Amastris_sp._GFDY01039857.1.p1
Cyphonia_clavata_GFEC01072213.1.p1
Cyphonia_clavata_GFEC01072214.1.p1
Stictocephala_bisonia_GCZK01029075.1.p1
Procyrta_sp._GFFD01037600.1.p1
Procyrta_sp._GFFD01040868.1.p1
Notocera_sp._GFFB01042487.1.p1
Membracis_tectigera_GFET01058677.1.p1
Lophyraspis_sp._GFEQ01031296.1.p1
Aetalion_reticulatum_GFDW01061011.1.p1
Lycoderes_burmeisteri_GFER01080073.1.p1
Lycoderes_burmeisteri_GFER01080074.1.p1
Centrotus_cornutus_GDFF01015164.1.p1
Llanquihuea_pilosa_GCWX01038011.1.p1
Llanquihuea_pilosa_GCWX01038012.1.p1
Idiocerus_rotundens_GEJG01018920.1.p1
Tinobregmus_viridescens_GEIL01013128.1.p1
Neocoelidia_tumidifrons_GEJC01043982.1.p1
Penthimia_sp._GFFF01051349.1.p1
Neoaliturus_tenellus_nte_C308178.p1
Agudus_sp._GFDX01080727.1.p1
Macrosteles_quadrilineatus_ref_MQ_DN33246_c0_g1_i1_2__frr
Agallia_constricta_GCWT01029690.1.p1
Ulopa_reticulata_GDE001019207.1.p1
Nionia_palmeri_GEKN01047408.1.p1
Hespenedra_chilensis_GCXG01008013.1.p1
Xestocephalus_desertorum_GELC01033665.1.p1
Macropsis_decisa_GEJE01017378.1.p1
Aphrodes_bicincta_GFDZ01148872.1.p1
Aphrodes_bicincta_GFDZ01116408.1.p1
Aphrodes_bicincta_GFDZ01116409.1.p1
Mapuchea_sp._GCXN01048506.1.p1
Mapuchea_sp._GCXN01048507.1.p1
MBK8907438.1
PJB70712.1
MAF68089.1
MAH06195.1
WP_070157063.1
WP_022672746.1
WP_167095179.1
MAB15174.1
WP_109919868.1
MBL4739070.1
0.2

*gh25*

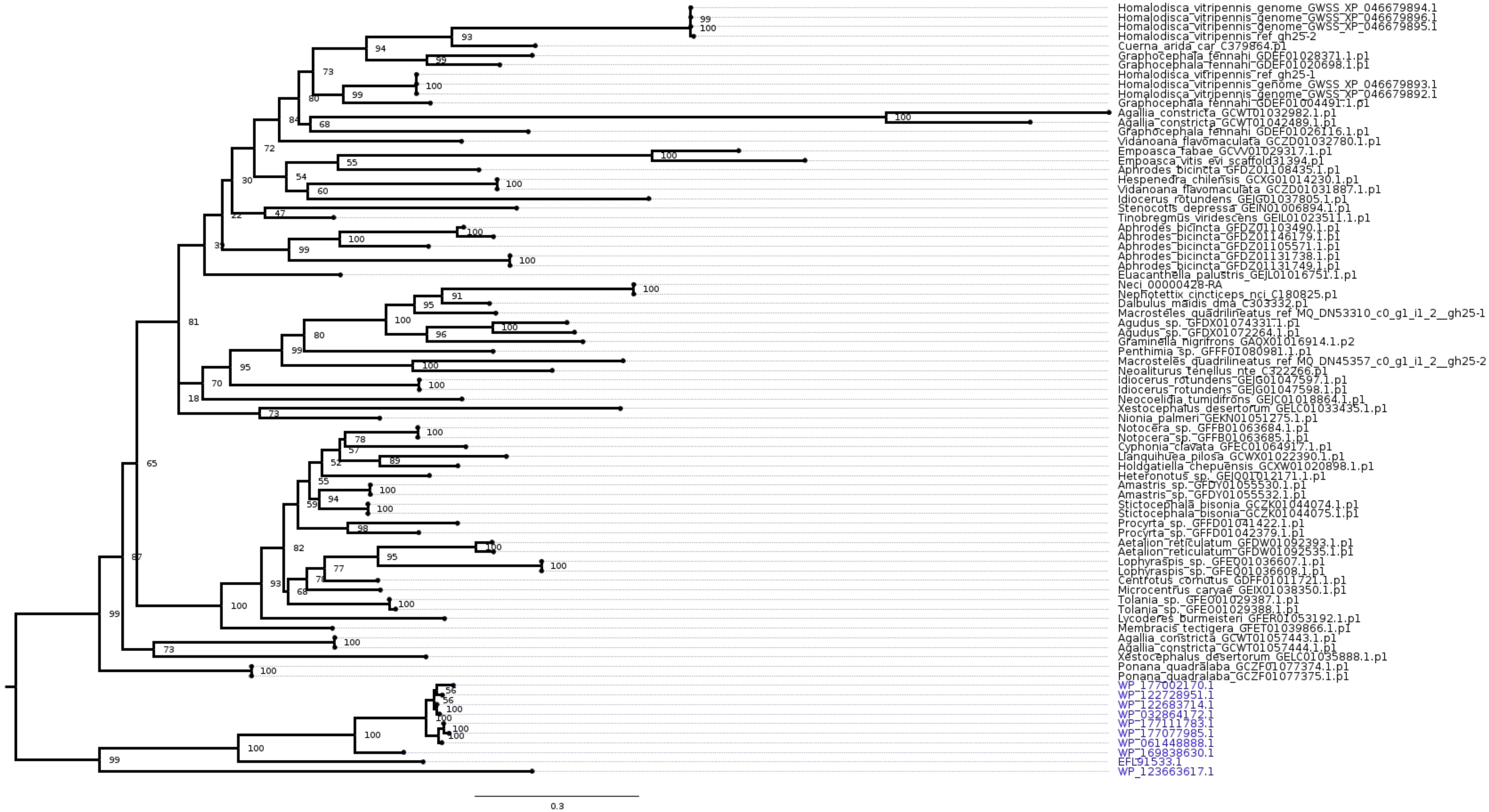

Homalodisca_vitripennis_genome_GWSS_XP_046679894.1
Homalodisca_vitripennis_genome_GWSS_XP_046679896.1
Homalodisca_vitripennis_genome_GWSS_XP_046679895.1
Homalodisca_vitripennis_ref_gh25-2
Cuerna_arida_car_C379864.p1
Graphocephala_fennahi_GDEF01028371.1.p1
Graphocephala_fennahi_GDEF01020698.1.p1
Homalodisca_vitripennis_ref_gh25-1
Homalodisca_vitripennis_genome_GWSS_XP_046679893.1
Homalodisca_vitripennis_genome_GWSS_XP_046679892.1
Graphocephala_fennahi_GDEF01004491.1.p1
Agallia_constricta_GCWT01032982.1.p1
Agallia_constricta_GCWT01042489.1.p1
Graphocephala_fennahi_GDEF01026116.1.p1
Vidanoana_flavomaculata_GCZD01032780.1.p1
Empoasca_fabae_GCVV01029317.1.p1
Empoasca_vitis_evi_scaffold31394.p1
Aphrodes_bicincta_GFDZ01108435.1.p1
Hespenedra_chilensis_GCXG01014230.1.p1
Vidanoana_flavomaculata_GCZD01031887.1.p1
Idiocerus_rotundens_GEJG01037805.1.p1
Stenocotis_depressa_GEIN01006894.1.p1
Tinobregmus_viridescens_GEIL01023511.1.p1
Aphrodes_bicincta_GFDZ01103490.1.p1
Aphrodes_bicincta_GFDZ01146179.1.p1
Aphrodes_bicincta_GFDZ01105571.1.p1
Aphrodes_bicincta_GFDZ01131738.1.p1
Aphrodes_bicincta_GFDZ01131749.1.p1
Euacanthella_palustris_GEJL01016751.1.p1
Neci_00000428-RA
Nephotettix_cincticeps_nci_C180825.p1
Dalbulus_maidis_dma_C303332.p1
Macrosteles_quadrilineatus_ref_MQ_DN53310_c0_g1_i1_2__gh25-1
Agudus_sp._GFDX01074331.1.p1
Agudus_sp._GFDX01072264.1.p1
Graminella_nigrifrons_GAQX01016914.1.p2
Penthimia_sp._GFFF01080981.1.p1
Macrosteles_quadrilineatus_ref_MQ_DN45357_c0_g1_i1_2__gh25-2
Neoaliturus_tenellus_nte_C322266.p1
Idiocerus_rotundens_GEJG01047597.1.p1
Idiocerus_rotundens_GEJG01047598.1.p1
Neocoelidia_tumidifrons_GEJC01018864.1.p1
Xestocephalus_desertorum_GELC01033435.1.p1
Nionia_palmeri_GEKN01051275.1.p1
Notocera_sp._GFFB01063684.1.p1
Notocera_sp._GFFB01063685.1.p1
Cyphonia_clavata_GFEC01064917.1.p1
Llanquihuea_pilosa_GCWX01022390.1.p1
Holdgatiella_chepuensis_GCXW01020898.1.p1
Heteronotus_sp._GEJQ01012171.1.p1
Amastris_sp._GFDY01055530.1.p1
Amastris_sp._GFDY01055532.1.p1
Stictocephala_bisonia_GCZK01044074.1.p1
Stictocephala_bisonia_GCZK01044075.1.p1
Procyrta_sp._GFFD01041422.1.p1
Procyrta_sp._GFFD01042379.1.p1
Aetalion_reticulatum_GFDW01092393.1.p1
Aetalion_reticulatum_GFDW01092535.1.p1
Lophyraspis_sp._GFEQ01036607.1.p1
Lophyraspis_sp._GFEQ01036608.1.p1
Centrotus_cornutus_GDFF01011721.1.p1
Microcentrus_caryae_GEIX01038350.1.p1
Tolania_sp._GFEO01029387.1.p1
Tolania_sp._GFEO01029388.1.p1
Lycoderes_burmeisteri_GFER01053192.1.p1
Membracis_tectigera_GFET01039866.1.p1
Agallia_constricta_GCWT01057443.1.p1
Agallia_constricta_GCWT01057444.1.p1
Xestocephalus_desertorum_GELC01035888.1.p1
Ponana_quadralaba_GCZF01077374.1.p1
Ponana_quadralaba_GCZF01077375.1.p1
WP_177002170.1
WP_122728951.1
WP_122683714.1
WP_032864172.1
WP_177111783.1
WP_177077985.1
WP_061448888.1
WP_169838630.1
EFL91533.1
WP_123663617.1
0.3

### *ileS*

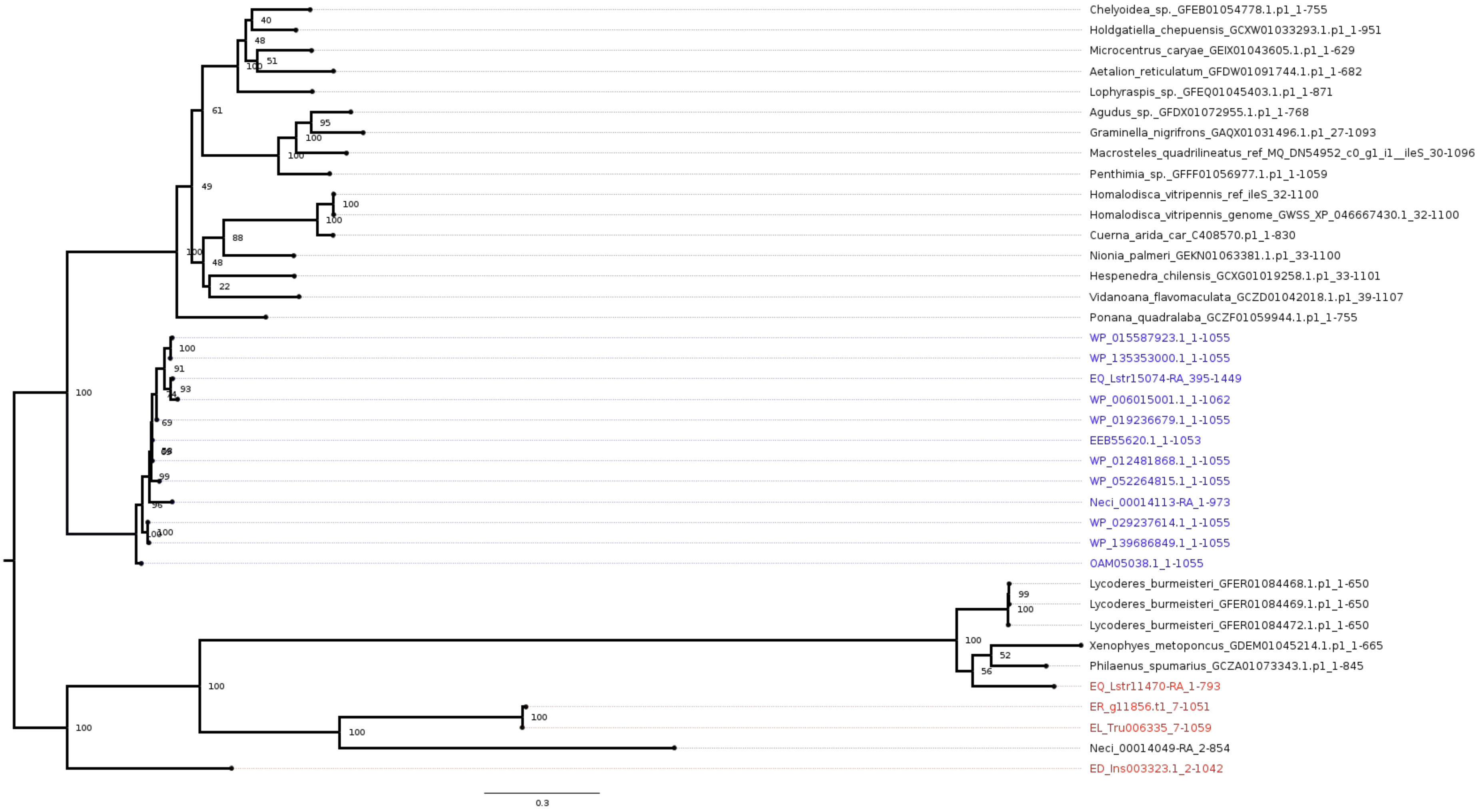
Chelyoidea_sp._GFEB01054778.1.p1_1-755
Holdgatiella_chepuensis_GCXW01033293.1.p1_1-951
Microcentrus_caryae_GEIX01043605.1.p1_1-629
Aetalion_reticulatum_GFDW01091744.1.p1_1-682
Lophyraspis_sp._GFEQ01045403.1.p1_1-871
Agudus_sp._GFDX01072955.1.p1_1-768
Graminella_nigrifrons_GAQX01031496.1.p1_27-1093
Macrosteles_quadrilineatus_ref_MQ_DN54952_c0_g1_i1__ileS_30-1096
Penthimia_sp._GFFF01056977.1.p1_1-1059
Homalodisca_vitripennis_ref_ileS_32-1100
Homalodisca_vitripennis_genome_GWSS_XP_046667430.1_32-1100
Cuerna_arida_car_C408570.p1_1-830
Nionia_palmeri_GEKN01063381.1.p1_33-1100
Hespenedra_chilensis_GCXG01019258.1.p1_33-1101
Vidanoana_flavomaculata_GCZD01042018.1.p1_39-1107
Ponana_quadralaba_GCZF01059944.1.p1_1-755
WP_015587923.1_1-1055
WP_135353000.1_1-1055
EQ_Lstr15074-RA_395-1449
WP_006015001.1_1-1062
WP_019236679.1_1-1055
EEB55620.1_1-1053
WP_012481868.1_1-1055
WP_052264815.1_1-1055
Neci_00014113-RA_1-973
WP_029237614.1_1-1055
WP_139686849.1_1-1055
OAM05038.1_1-1055
Lycoderes_burmeisteri_GFER01084468.1.p1_1-650
Lycoderes_burmeisteri_GFER01084469.1.p1_1-650
Lycoderes_burmeisteri_GFER01084472.1.p1_1-650
Xenophyes_metoponcus_GDEM01045214.1.p1_1-665
Philaenus_spumarius_GCZA01073343.1.p1_1-845
EQ_Lstr11470-RA_1-793
ER_g11856.t1_7-1051
EL_Tru006335_7-1059
Neci_00014049-RA_2-854
ED_Ins003323.1_2-1042
0.3

### *pel*

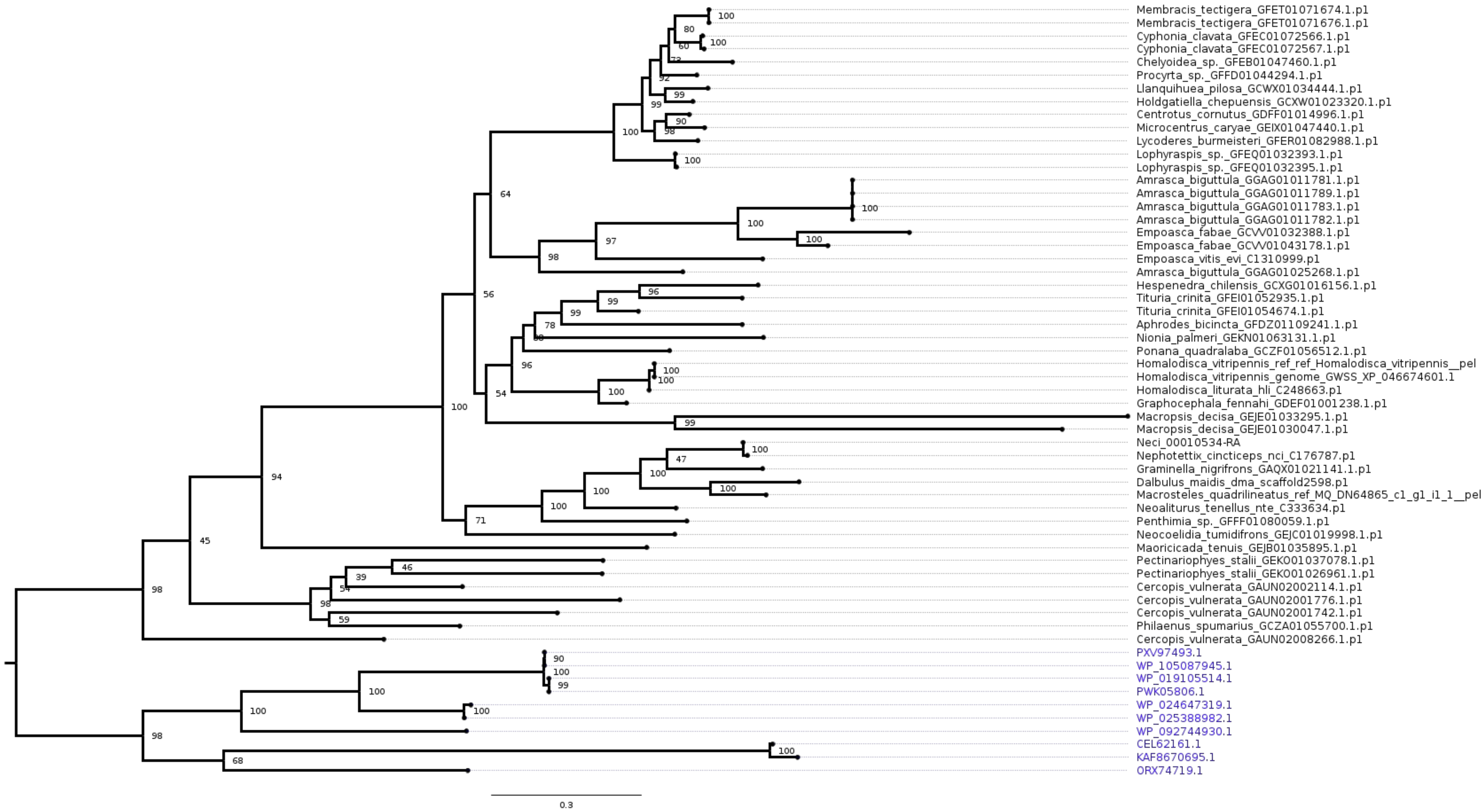

Membracis_tectigera_GFET01071674.1.p1
Membracis_tectigera_GFET01071676.1.p1
Cyphonia_clavata_GFEC01072566.1.p1
Cyphonia_clavata_GFEC01072567.1.p1
Chelyoidea_sp._GFEB01047460.1.p1
Procyrta_sp._GFFD01044294.1.p1
Llanquihuea_pilosa_GCWX01034444.1.p1
Holdgatiella_chepuensis_GCXW01023320.1.p1
Centrotus_cornutus_GDFF01014996.1.p1
Microcentrus_caryae_GEIX01047440.1.p1
Lycoderes_burmeisteri_GFER01082988.1.p1
Lophyraspis_sp._GFEQ01032393.1.p1
Lophyraspis_sp._GFEQ01032395.1.p1
Amrasca_biguttula_GGAG01011781.1.p1
Amrasca_biguttula_GGAG01011789.1.p1
Amrasca_biguttula_GGAG01011783.1.p1
Amrasca_biguttula_GGAG01011782.1.p1
Empoasca_fabae_GCVV01032388.1.p1
Empoasca_fabae_GCVV01043178.1.p1
Empoasca_vitis_evi_C1310999.p1
Amrasca_biguttula_GGAG01025268.1.p1
Hespenedra_chilensis_GCXG01016156.1.p1
Tituria_crinita_GFEI01052935.1.p1
Tituria_crinita_GFEI01054674.1.p1
Aphrodes_bicincta_GFDZ01109241.1.p1
Nionia_palmeri_GEKN01063131.1.p1
Ponana_quadralaba_GCZF01056512.1.p1
Homalodisca_vitripennis_ref_ref_Homalodisca_vitripennis__pel
Homalodisca_vitripennis_genome_GWSS_XP_046674601.1
Homalodisca_liturata_hli_C248663.p1
Graphocephala_fennahi_GDEF01001238.1.p1
Macropsis_decisa_GEJE01033295.1.p1
Macropsis_decisa_GEJE01030047.1.p1
Neci_00010534-RA
Nephotettix_cincticeps_nci_C176787.p1
Graminella_nigrifrons_GAQX01021141.1.p1
Dalbulus_maidis_dma_scaffold2598.p1
Macrosteles_quadrilineatus_ref_MQ_DN64865_c1_g1_i1_1__pel
Neoaliturus_tenellus_nte_C333634.p1
Penthimia_sp._GFFF01080059.1.p1
Neocoelidia_tumidifrons_GEJC01019998.1.p1
Maoricicada_tenuis_GEJB01035895.1.p1
Pectinariophyes_stalii_GEK001037078.1.p1
Pectinariophyes_stalii_GEK001026961.1.p1
Cercopis_vulnerata_GAUN02002114.1.p1
Cercopis_vulnerata_GAUN02001776.1.p1
Cercopis_vulnerata_GAUN02001742.1.p1
Philaenus_spumarius_GCZA01055700.1.p1
Cercopis_vulnerata_GAUN02008266.1.p1
PXV97493.1
WP_105087945.1
WP_019105514.1
PWK05806.1
WP_024647319.1
WP_025388982.1
WP_092744930.1
CEL62161.1
KAF8670695.1
ORX74719.1
0.3

### *per*

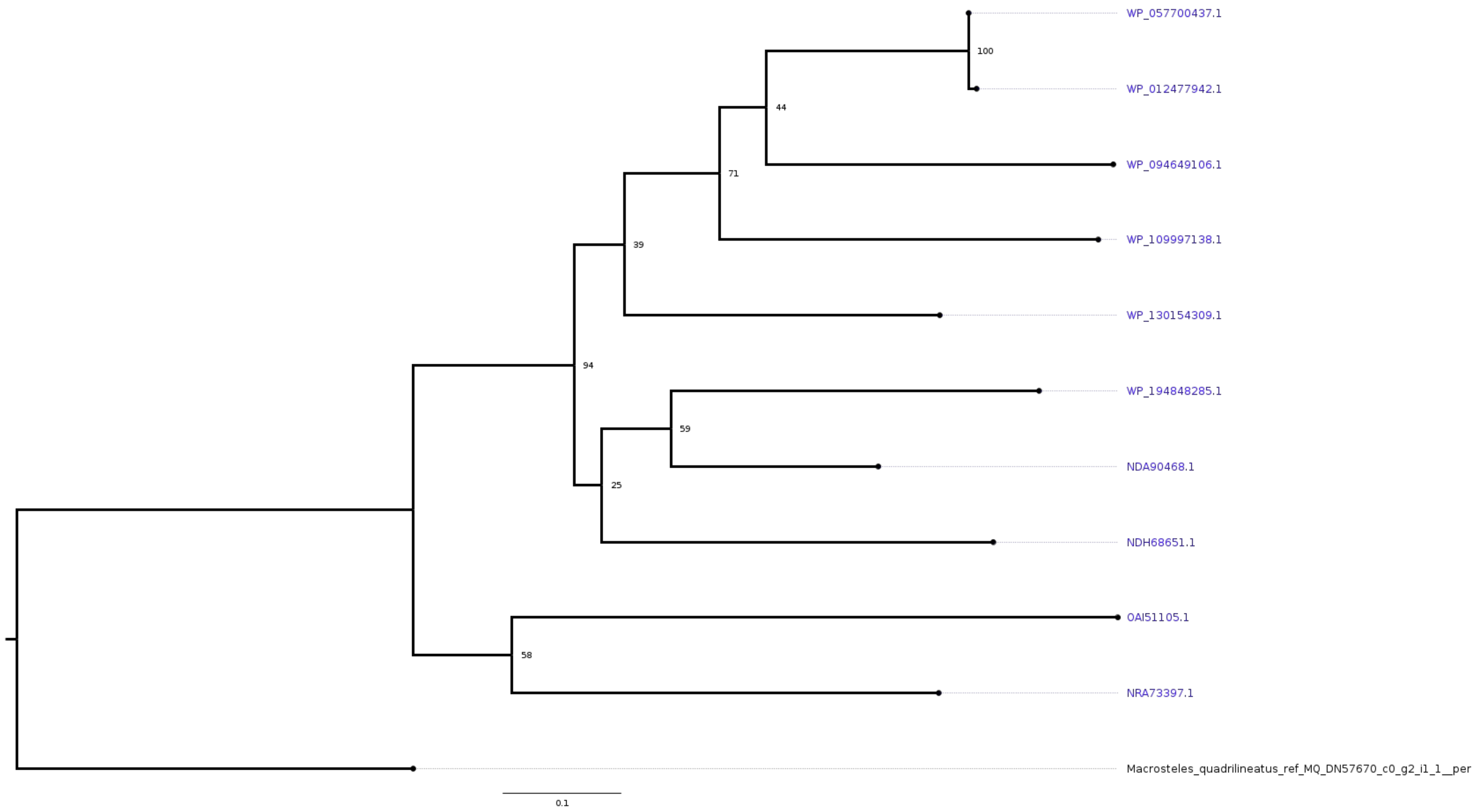

WP_057700437.1
100
WP_012477942.1
44
WP_094649106.1
71
WP_109997138.1
39
WP_130154309.1
94
WP_194848285.1
59
NDA90468.1
25
NDH68651.1
OAI51105.1
58
NRA73397.1
Macrosteles_quadrilineatus_ref_MQ_DN57670_c0_g2_i1_1__per
0.1

#### *plc*

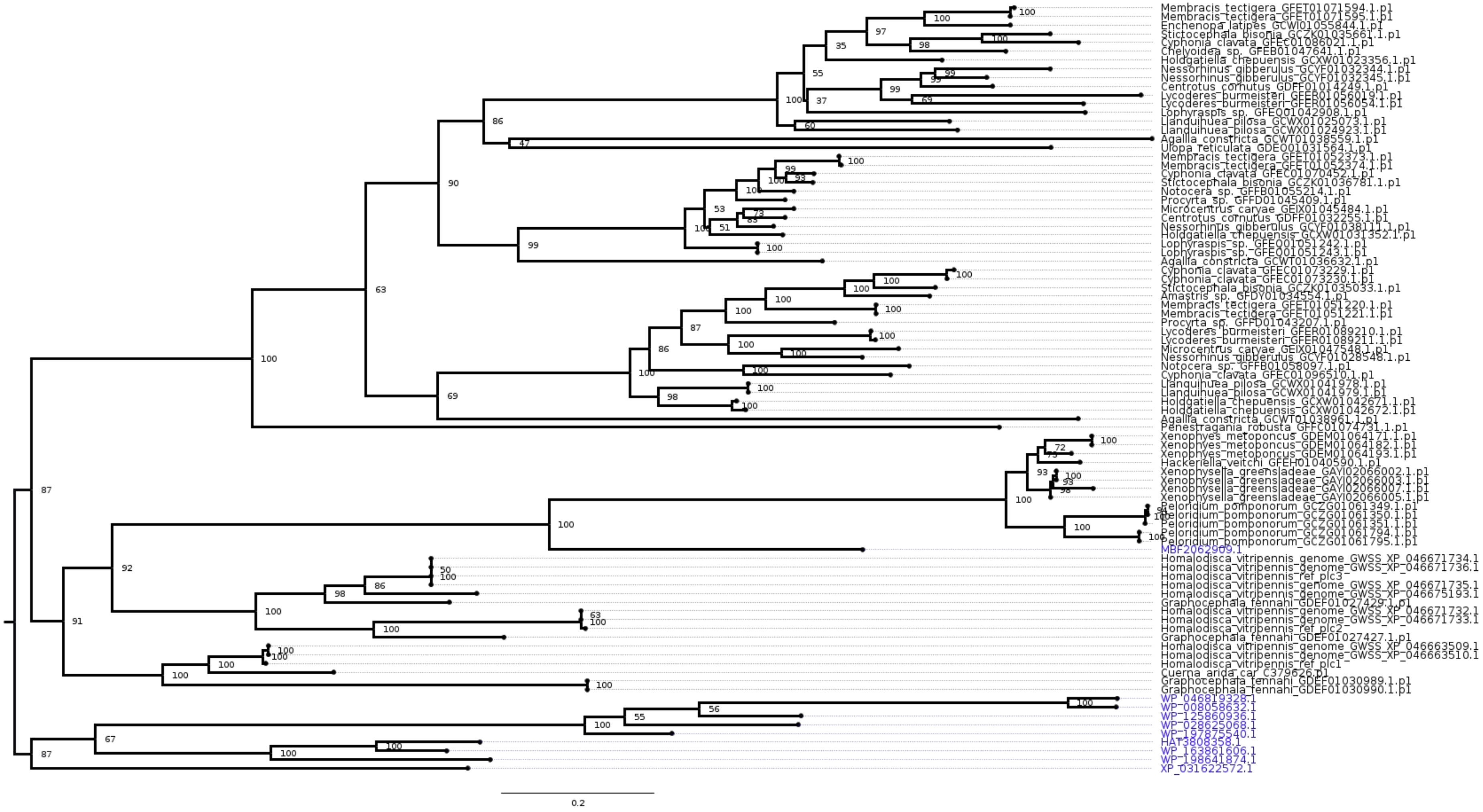

Membracis_tectigera_GFET01071594.1.p1
Membracis_tectigera_GFET01071595.1.p1
Enchenopa_latipes_GCWI01055844.1.p1
Stictocephala_bisonia_GCZK01035661.1.p1
Cyphonia_clavata_GFEC01086021.1.p1
Chelyoidea_sp._GFEB01047641.1.p1
Holdgatiella_chepuensis_GCXW01023356.1.p1
Nessorhinus_gibberulus_GCYF01032344.1.p1
Nessorhinus_gibberulus_GCYF01032345.1.p1
Centrotus_cornutus_GDFF01014249.1.p1
Lycoderes_burmeisteri_GFER01056019.1.p1
Lycoderes_burmeisteri_GFER01056054.1.p1
Lophyraspis_sp._GFEQ01042908.1.p1
Llanquihuea_pilosa_GCWX01025073.1.p1
Llanquihuea_pilosa_GCWX01024923.1.p1
Agallia_constricta_GCWT01038559.1.p1
Ulopa_reticulata_GDEO01031564.1.p1
Membracis_tectigera_GFET01052373.1.p1
Membracis_tectigera_GFET01052374.1.p1
Cyphonia_clavata_GFEC01070452.1.p1
Stictocephala_bisonia_GCZK01036781.1.p1
Notocera_sp._GFFB01055214.1.p1
Procyrta_sp._GFFD01045409.1.p1
Microcentrus_caryae_GEIX01045484.1.p1
Centrotus_cornutus_GDFF01032255.1.p1
Nessorhinus_gibberulus_GCYF01038111.1.p1
Holdgatiella_chepuensis_GCXW01031352.1.p1
Lophyraspis_sp._GFEQ01051242.1.p1
Lophyraspis_sp._GFEQ01051243.1.p1
Agallia_constricta_GCWT01036632.1.p1
Cyphonia_clavata_GFEC01073229.1.p1
Cyphonia_clavata_GFEC01073230.1.p1
Stictocephala_bisonia_GCZK01035033.1.p1
Amastris_sp._GFDY01034554.1.p1
Membracis_tectigera_GFET01051220.1.p1
Membracis_tectigera_GFET01051221.1.p1
Procyrta_sp._GFFD01043207.1.p1
Lycoderes_burmeisteri_GFER01089210.1.p1
Lycoderes_burmeisteri_GFER01089211.1.p1
Microcentrus_caryae_GEIX01047548.1.p1
Nessorhinus_gibberulus_GCYF01028548.1.p1
Notocera_sp._GFFB01058097.1.p1
Cyphonia_clavata_GFEC01096510.1.p1
Llanquihuea_pilosa_GCWX01041978.1.p1
Llanquihuea_pilosa_GCWX01041979.1.p1
Holdgatiella_chepuensis_GCXW01042671.1.p1
Holdgatiella_chepuensis_GCXW01042672.1.p1
Agallia_constricta_GCWT01038961.1.p1
Penestragania_robusta_GFFC01074731.1.p1
Xenophyes_metoponcus_GDEM01064171.1.p1
Xenophyes_metoponcus_GDEM01064182.1.p1
Xenophyes_metoponcus_GDEM01064193.1.p1
Hackeriella_veitchi_GFEH01040590.1.p1
Xenophysella_greensladeae_GAYI02066002.1.p1
Xenophysella_greensladeae_GAYI02066003.1.p1
Xenophysella_greensladeae_GAYI02066007.1.p1
Xenophysella_greensladeae_GAYI02066005.1.p1
Peloridium_pomponorum_GCZG01061349.1.p1
Peloridium_pomponorum_GCZG01061350.1.p1
Peloridium_pomponorum_GCZG01061351.1.p1
Peloridium_pomponorum_GCZG01061794.1.p1
Peloridium_pomponorum_GCZG01061795.1.p1
MBF2062909.1
Homalodisca_vitripennis_genome_GWSS_XP_046671734.1
Homalodisca_vitripennis_genome_GWSS_XP_046671736.1
Homalodisca_vitripennis_ref_plc3
Homalodisca_vitripennis_genome_GWSS_XP_046671735.1
Homalodisca_vitripennis_genome_GWSS_XP_046675193.1
Graphocephala_fennahi_GDEF01027429.1.p1
Homalodisca_vitripennis_genome_GWSS_XP_046671732.1
Homalodisca_vitripennis_genome_GWSS_XP_046671733.1
Homalodisca_vitripennis_ref_plc2
Graphocephala_fennahi_GDEF01027427.1.p1
Homalodisca_vitripennis_genome_GWSS_XP_046663509.1
Homalodisca_vitripennis_genome_GWSS_XP_046663510.1
Homalodisca_vitripennis_ref_plc1
Cuerna_arida_car_C379626.p1
Graphocephala_fennahi_GDEF01030989.1.p1
Graphocephala_fennahi_GDEF01030990.1.p1
WP_046819328.1
WP_008058632.1
WP_125860936.1
WP_028625068.1
WP_197875540.1
HAT3808358.1
WP_163861606.1
WP_198641874.1
XP_031622572.1
0.2

*ribD*

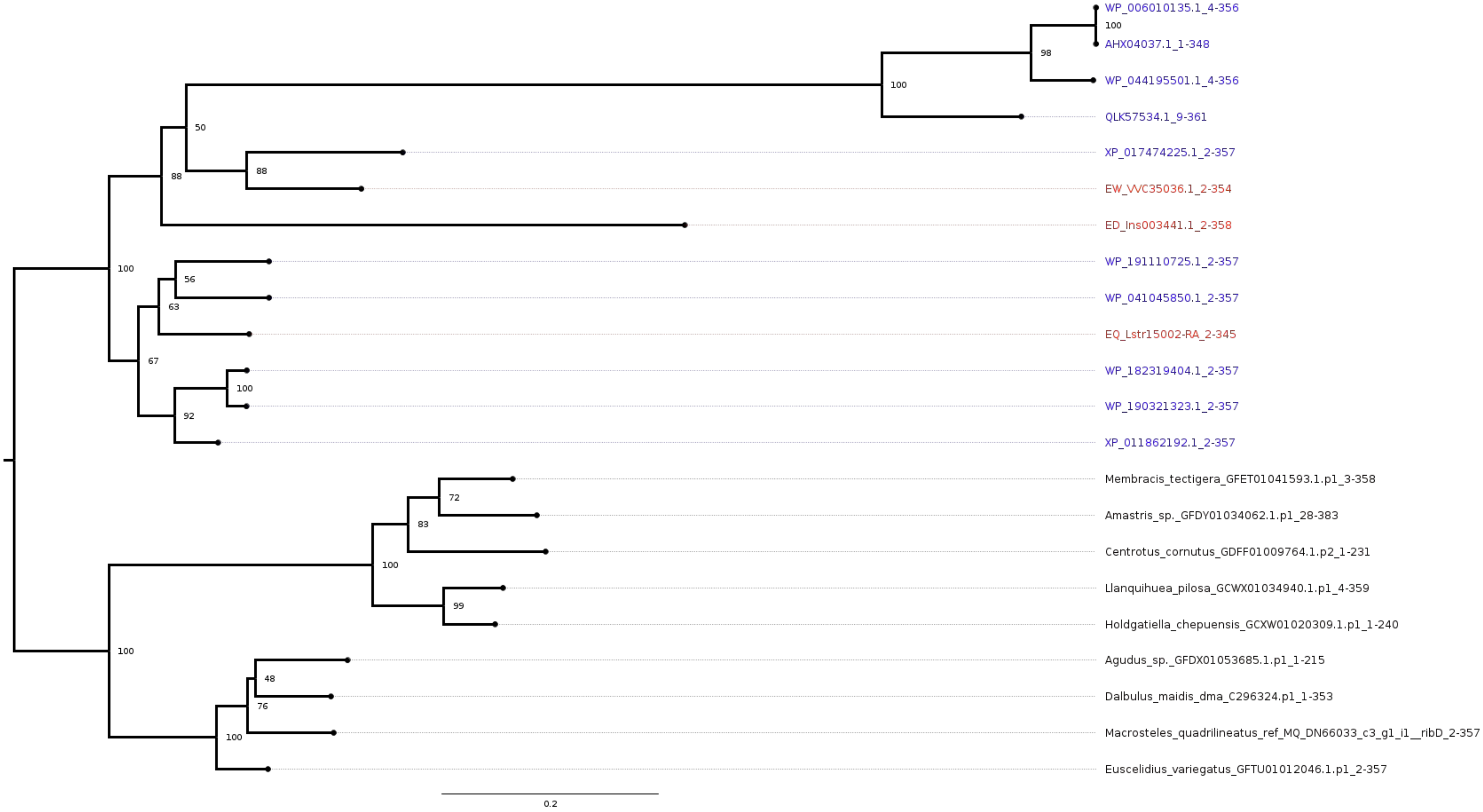
WP_006010135.1_4-356
100
AHX04037.1_1-348
98
WP_044195501.1_4-356
100
QLK57534.1_9-361
50
XP_017474225.1_2-357
88
EW_VVC35036.1_2-354
88
ED_Ins003441.1_2-358
100
WP_191110725.1_2-357
56
WP_041045850.1_2-357
63
EQ_Lstr15002-RA_2-345
67
WP_182319404.1_2-357
100
WP_190321323.1_2-357
92
XP_011862192.1_2-357
Membracis_tectigera_GFET01041593.1.p1_3-358
72
Amastris_sp._GFDY01034062.1.p1_28-383
83
Centrotus_cornutus_GDFF01009764.1.p2_1-231
100
Llanquihuea_pilosa_GCWX01034940.1.p1_4-359
99
Holdgatiella_chepuensis_GCXW01020309.1.p1_1-240
100
Agudus_sp._GFDX01053685.1.p1_1-215
48
Dalbulus_maidis_dma_C296324.p1_1-353
76
Macrosteles_quadrilineatus_ref_MQ_DN66033_c3_g1_i1__ribD_2-357
100
Euscelidius_variegatus_GFTU01012046.1.p1_2-357
0.2

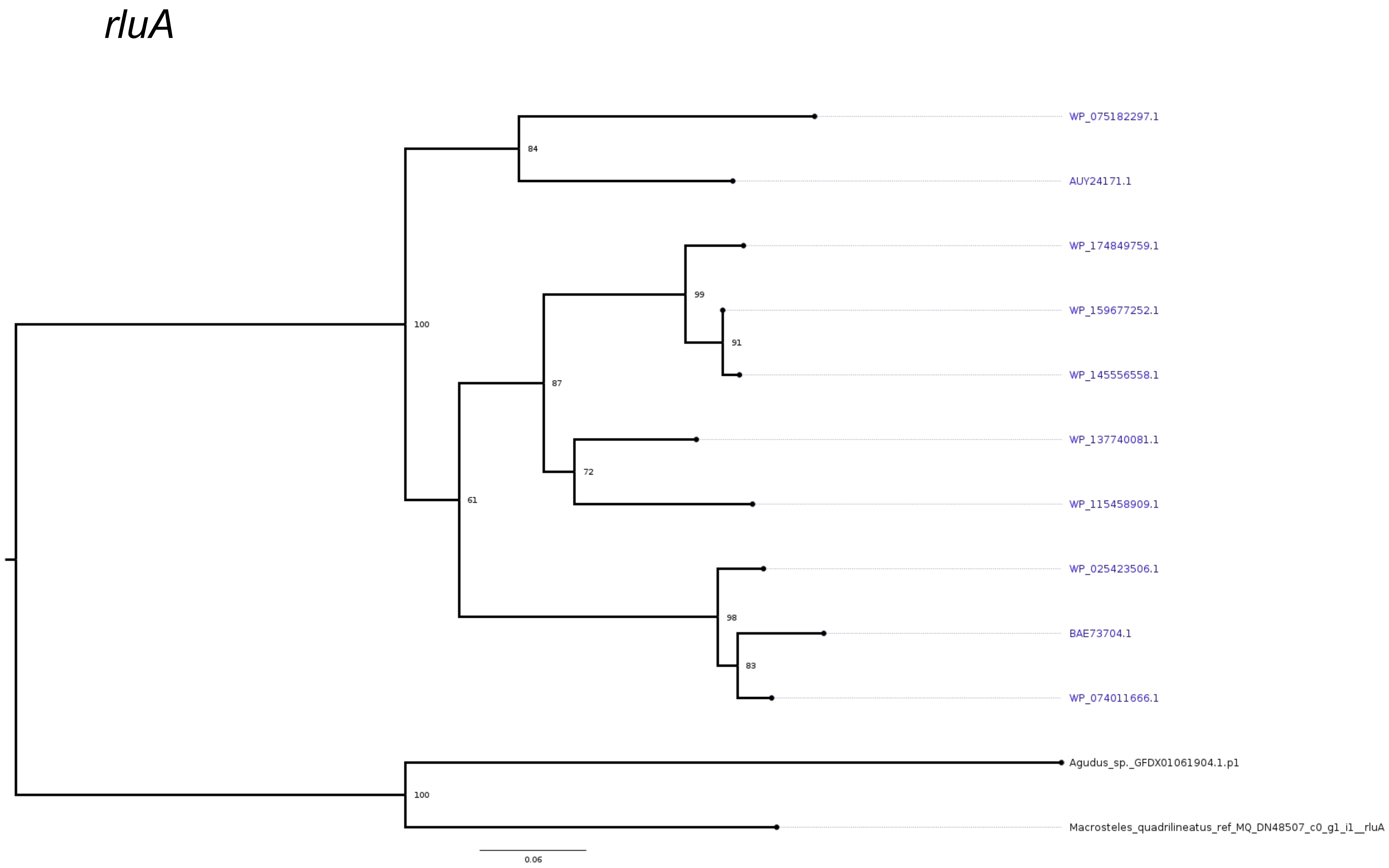
rluA
WP_075182297.1
AUY24171.1
WP_174849759.1
WP_159677252.1
WP_145556558.1
WP_137740081.1
WP_115458909.1
WP_025423506.1
BAE73704.1
WP_074011666.1
Agudus_sp._GFDX01061904.1.p1
Macrosteles_quadrilineatus_ref_MQ_DN48507_c0_g1_i1__rluA
84
100
99
91
87
72
61
98
83
100
0.06

*rnc*

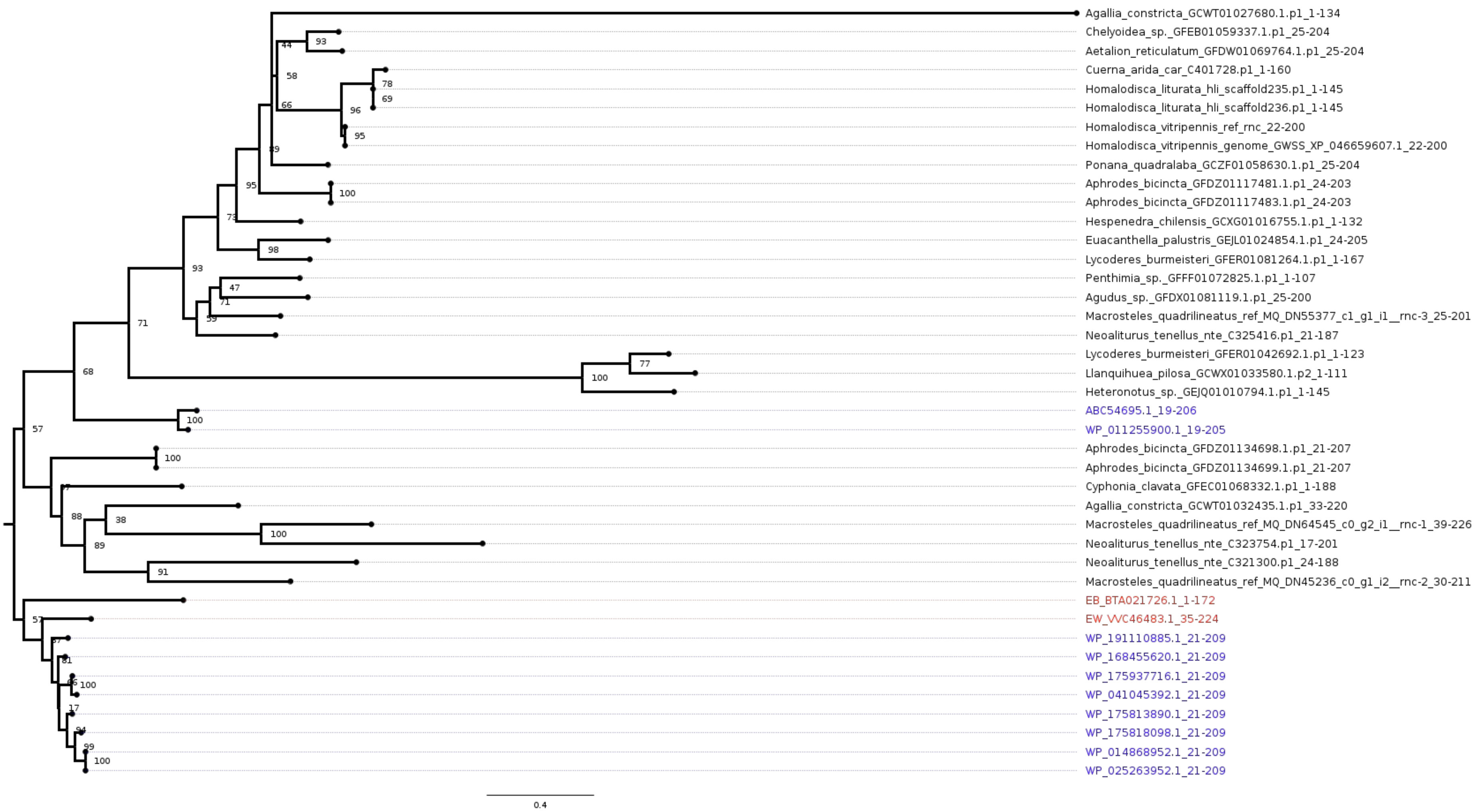
Agallia_constricta_GCWT01027680.1.p1_1-134
Chelyoidea_sp._GFEB01059337.1.p1_25-204
Aetalion_reticulatum_GFDW01069764.1.p1_25-204
Cuerna_arida_car_C401728.p1_1-160
Homalodisca_liturata_hli_scaffold235.p1_1-145
Homalodisca_liturata_hli_scaffold236.p1_1-145
Homalodisca_vitripennis_ref_rnc_22-200
Homalodisca_vitripennis_genome_GWSS_XP_046659607.1_22-200
Ponana_quadralaba_GCZF01058630.1.p1_25-204
Aphrodes_bicincta_GFDZ01117481.1.p1_24-203
Aphrodes_bicincta_GFDZ01117483.1.p1_24-203
Hespenedra_chilensis_GCXG01016755.1.p1_1-132
Euacanthella_palustris_GEJL01024854.1.p1_24-205
Lycoderes_burmeisteri_GFER01081264.1.p1_1-167
Penthimia_sp._GFFF01072825.1.p1_1-107
Agudus_sp._GFDX01081119.1.p1_25-200
Macrosteles_quadrilineatus_ref_MQ_DN55377_c1_g1_i1__rnc-3_25-201
Neoaliturus_tenellus_nte_C325416.p1_21-187
Lycoderes_burmeisteri_GFER01042692.1.p1_1-123
Llanquihuea_pilosa_GCWX01033580.1.p2_1-111
Heteronotus_sp._GEJQ01010794.1.p1_1-145
ABC54695.1_19-206
WP_011255900.1_19-205
Aphrodes_bicincta_GFDZ01134698.1.p1_21-207
Aphrodes_bicincta_GFDZ01134699.1.p1_21-207
Cyphonia_clavata_GFEC01068332.1.p1_1-188
Agallia_constricta_GCWT01032435.1.p1_33-220
Macrosteles_quadrilineatus_ref_MQ_DN64545_c0_g2_i1__rnc-1_39-226
Neoaliturus_tenellus_nte_C323754.p1_17-201
Neoaliturus_tenellus_nte_C321300.p1_24-188
Macrosteles_quadrilineatus_ref_MQ_DN45236_c0_g1_i2__rnc-2_30-211
EB_BTA021726.1_1-172
EW_VVC46483.1_35-224
WP_191110885.1_21-209
WP_168455620.1_21-209
WP_175937716.1_21-209
WP_041045392.1_21-209
WP_175813890.1_21-209
WP_175818098.1_21-209
WP_014868952.1_21-209
WP_025263952.1_21-209
0.4

*tmk*

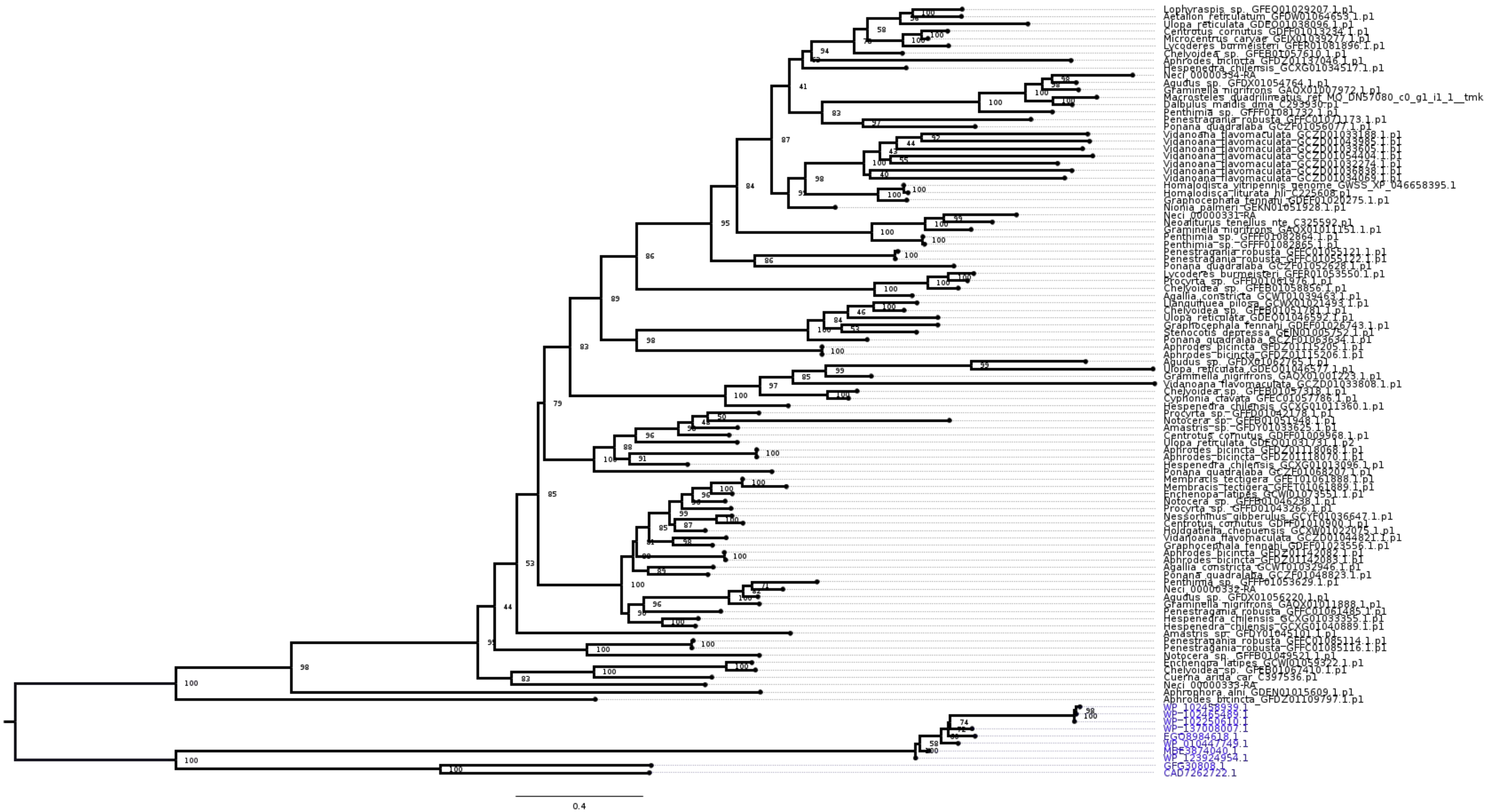
Lophyraspis_sp._GFEQ01029207.1.p1
Aetalion_reticulatum_GFDW01064653.1.p1
Ulopa_reticulata_GDEO01038096.1.p1
Centrotus_cornutus_GDFF01013234.1.p1
Microcentrus_caryae_GEIX01039277.1.p1
Lycoderes_burmeisteri_GFER01081896.1.p1
Chelyoidea_sp._GFEB01057610.1.p1
Aphrodes_bicincta_GFDZ01137046.1.p1
Hespenedra_chilensis_GCXG01034517.1.p1
Neci_00000334-RA
Agudus_sp._GFDX01054764.1.p1
Graminella_nigrifrons_GAQX01007972.1.p1
Macrosteles_quadrilineatus_ret_MQ_DN57080_c0_g1_i1_1__tmk
Dalbulus_maidis_dma_C293930.p1
Penthimia_sp._GFFF01081732.1.p1
Penestragania_robusta_GFFC01071173.1.p1
Ponana_quadralaba_GCZF01056077.1.p1
Vidanoana_flavomaculata_GCZD01033188.1.p1
Vidanoana_flavomaculata_GCZD01043985.1.p1
Vidanoana_flavomaculata_GCZD01033605.1.p1
Vidanoana_flavomaculata_GCZD01054404.1.p1
Vidanoana_flavomaculata_GCZD01032274.1.p1
Vidanoana_flavomaculata_GCZD01036838.1.p1
Vidanoana_flavomaculata_GCZD01034069.1.p1
Homalodisca_vitripennis_genome_GWSS_XP_046658395.1
Homalodisca_liturata_hli_C225608.p1
Graphocephala_fennahi_GDEF01020275.1.p1
Nionia_palmeri_GEKN01051928.1.p1
Neci_00000331-RA
Neoaliturus_tenellus_nte_C325592.p1
Graminella_nigrifrons_GAQX01011151.1.p1
Penthimia_sp._GFFF01082864.1.p1
Penthimia_sp._GFFF01082865.1.p1
Penestragania_robusta_GFFC01055121.1.p1
Penestragania_robusta_GFFC01055122.1.p1
Ponana_quadralaba_GCZF01052628.1.p1
Lycoderes_burmeisteri_GFER01053550.1.p1
Procyrta_sp._GFFD01061976.1.p1
Chelyoidea_sp._GFEB01058856.1.p1
Agallia_constricta_GCWT01039463.1.p1
Llanquihuea_pilosa_GCWX01021493.1.p1
Chelyoidea_sp._GFEB01051781.1.p1
Ulopa_reticulata_GDEO01046592.1.p1
Graphocephala_fennahi_GDEF01026743.1.p1
Stenocotis_depressa_GEIN01005752.1.p1
Ponana_quadralaba_GCZF01063634.1.p1
Aphrodes_bicincta_GFDZ01115205.1.p1
Aphrodes_bicincta_GFDZ01115206.1.p1
Agudus_sp._GFDX01062765.1.p1
Ulopa_reticulata_GDEO01046577.1.p1
Graminella_nigrifrons_GAQX01001223.1.p1
Vidanoana_flavomaculata_GCZD01033808.1.p1
Chelyoidea_sp._GFEB01057318.1.p1
Cyphonia_clavata_GFEC01057786.1.p1
Hespenedra_chilensis_GCXG01011360.1.p1
Procyrta_sp._GFFD01042178.1.p1
Notocera_sp._GFFB01051948.1.p1
Amastris_sp._GFDY01033625.1.p1
Centrotus_cornutus_GDFF01009968.1.p1
Ulopa_reticulata_GDEO01031731.1.p2
Aphrodes_bicincta_GFDZ01118068.1.p1
Aphrodes_bicincta_GFDZ01118070.1.p1
Hespenedra_chilensis_GCXG01013096.1.p1
Ponana_quadralaba_GCZF01068207.1.p1
Membracis_tectigera_GFET01061888.1.p1
Membracis_tectigera_GFET01061889.1.p1
Enchenopa_latipes_GCWI01073551.1.p1
Notocera_sp._GFFB01046238.1.p1
Procyrta_sp._GFFD01043266.1.p1
Nessorhinus_gibberulus_GCYF01036647.1.p1
Centrotus_cornutus_GDFF01010900.1.p1
Holdgatiella_chepuensis_GCXW01022075.1.p1
Vidanoana_flavomaculata_GCZD01044821.1.p1
Graphocephala_fennahi_GDEF01023556.1.p1
Aphrodes_bicincta_GFDZ01142082.1.p1
Aphrodes_bicincta_GFDZ01142083.1.p1
Agallia_constricta_GCWT01032946.1.p1
Ponana_quadralaba_GCZF01048823.1.p1
Penthimia_sp._GFFF01053629.1.p1
Neci_00000332-RA
Agudus_sp._GFDX01056220.1.p1
Graminella_nigrifrons_GAQX01011888.1.p1
Penestragania_robusta_GFFC01061485.1.p1
Hespenedra_chilensis_GCXG01033355.1.p1
Hespenedra_chilensis_GCXG01040889.1.p1
Amastris_sp._GFDY01045101.1.p1
Penestragania_robusta_GFFC01085114.1.p1
Penestragania_robusta_GFFC01085116.1.p1
Notocera_sp._GFFB01049521.1.p1
Enchenopa_latipes_GCWI01059322.1.p1
Chelyoidea_sp._GFEB01067410.1.p1
Cuerna_arida_car_C397536.p1
Neci_00000333-RA
Aphrophora_alni_GDEN01015609.1.p1
Aphrodes_bicincta_GFDZ01109797.1.p1
WP_102458939.1
WP_102465489.1
WP_102250610.1
WP_137008007.1
EGO8984618.1
WP_010447749.1
MBE3874040.1
WP_123924954.1
GFG30808.1
CAD7262722.1
0.4

*yebC*

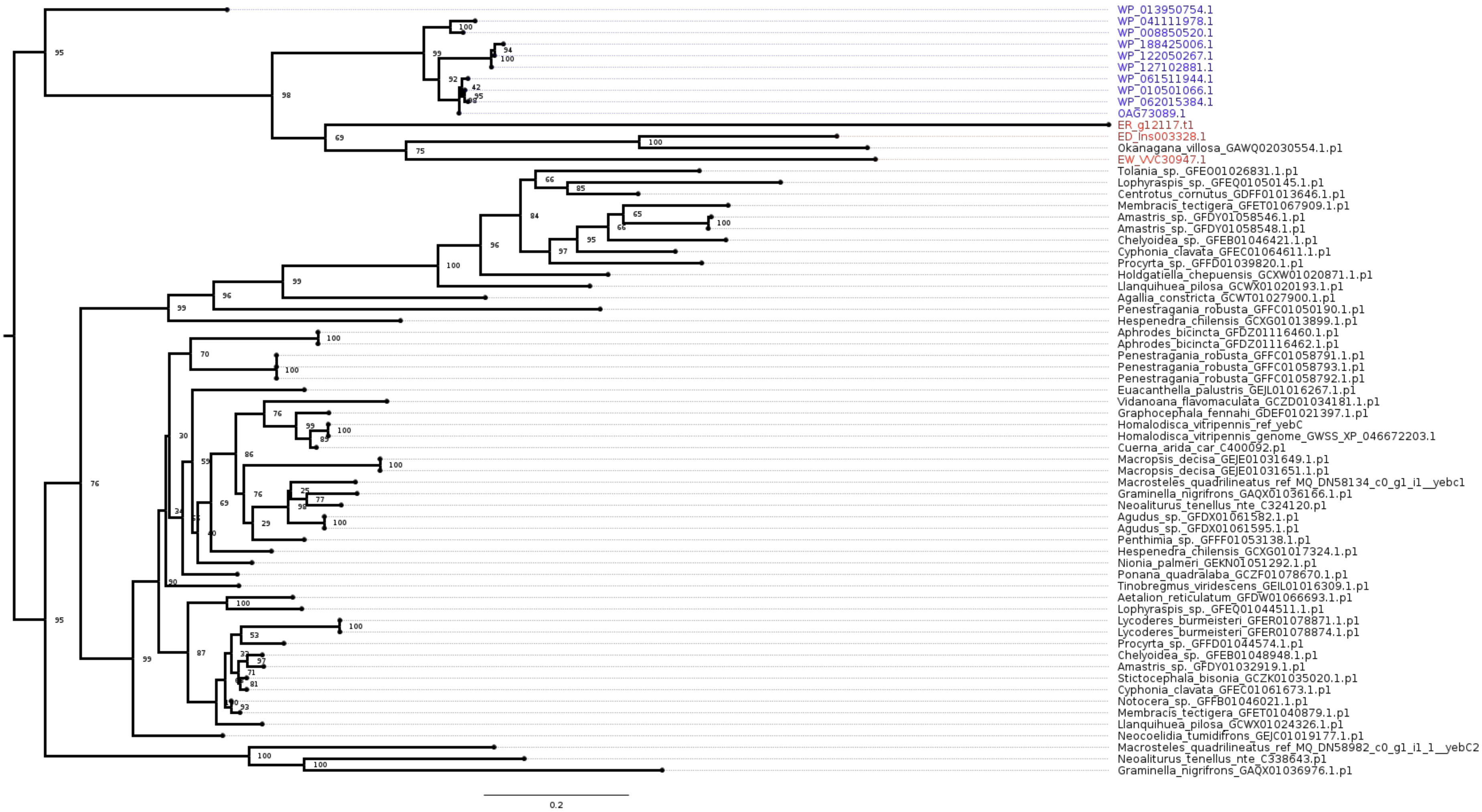
WP_013950754.1
WP_041111978.1
WP_008850520.1
WP_188425006.1
WP_122050267.1
WP_127102881.1
WP_061511944.1
WP_010501066.1
WP_062015384.1
OAG73089.1
ER_g12117.t1
ED_Ins003328.1
Okanagana_villosa_GAWQ02030554.1.p1
EW_VVC30947.1
Tolania_sp._GFEO01026831.1.p1
Lophyraspis_sp._GFEQ01050145.1.p1
Centrotus_cornutus_GDFF01013646.1.p1
Membracis_tectigera_GFET01067909.1.p1
Amastris_sp._GFDY01058546.1.p1
Amastris_sp._GFDY01058548.1.p1
Chelyoidea_sp._GFEB01046421.1.p1
Cyphonia_clavata_GFEC01064611.1.p1
Procyrta_sp._GFFD01039820.1.p1
Holdgatiella_chepuensis_GCXW01020871.1.p1
Llanquihuea_pilosa_GCWX01020193.1.p1
Agallia_constricta_GCWT01027900.1.p1
Penestragania_robusta_GFFC01050190.1.p1
Hespenedra_chilensis_GCXG01013899.1.p1
Aphrodes_bicincta_GFDZ01116460.1.p1
Aphrodes_bicincta_GFDZ01116462.1.p1
Penestragania_robusta_GFFC01058791.1.p1
Penestragania_robusta_GFFC01058793.1.p1
Penestragania_robusta_GFFC01058792.1.p1
Euacanthella_palustris_GEJL01016267.1.p1
Vidanoana_flavomaculata_GCZD01034181.1.p1
Graphocephala_fennahi_GDEF01021397.1.p1
Homalodisca_vitripennis_ref_yebC
Homalodisca_vitripennis_genome_GWSS_XP_046672203.1
Cuerna_arida_car_C400092.p1
Macropsis_decisa_GEJE01031649.1.p1
Macropsis_decisa_GEJE01031651.1.p1
Macrosteles_quadrilineatus_ref_MQ_DN58134_c0_g1_i1__yebc1
Graminella_nigrifrons_GAQX01036166.1.p1
Neoaliturus_tenellus_nte_C324120.p1
Agudus_sp._GFDX01061582.1.p1
Agudus_sp._GFDX01061595.1.p1
Penthimia_sp._GFFF01053138.1.p1
Hespenedra_chilensis_GCXG01017324.1.p1
Nionia_palmeri_GEKN01051292.1.p1
Ponana_quadralaba_GCZF01078670.1.p1
Tinobregmus_viridescens_GEIL01016309.1.p1
Aetalion_reticulatum_GFDW01066693.1.p1
Lophyraspis_sp._GFEQ01044511.1.p1
Lycoderes_burmeisteri_GFER01078871.1.p1
Lycoderes_burmeisteri_GFER01078874.1.p1
Procyrta_sp._GFFD01044574.1.p1
Chelyoidea_sp._GFEB01048948.1.p1
Amastris_sp._GFDY01032919.1.p1
Stictocephala_bisonia_GCZK01035020.1.p1
Cyphonia_clavata_GFEC01061673.1.p1
Notocera_sp._GFFB01046021.1.p1
Membracis_tectigera_GFET01040879.1.p1
Llanquihuea_pilosa_GCWX01024326.1.p1
Neocoelidia_tumidifrons_GEJC01019177.1.p1
Macrosteles_quadrilineatus_ref_MQ_DN58982_c0_g1_i1_1__yebC2
Neoaliturus_tenellus_nte_C338643.p1
Graminella_nigrifrons_GAQX01036976.1.p1
0.2
